# The contribution of gene flow, selection, and genetic drift to five thousand years of human allele frequency change

**DOI:** 10.1101/2023.07.11.548607

**Authors:** Alexis Simon, Graham Coop

## Abstract

Genomic time series from experimental evolution studies and ancient DNA datasets offer us a chance to directly observe the interplay of various evolutionary forces. We show how the genome-wide variance in allele frequency change between two time points can be decomposed into the contributions of gene flow, genetic drift, and linked selection. In closed populations, the contribution of linked selection is identifiable because it creates covariances between time intervals, and genetic drift does not. However, repeated gene flow between populations can also produce directionality in allele frequency change, creating covariances. We show how to accurately separate the fraction of variance in allele frequency change due to admixture and linked selection in a population receiving gene flow. We use two human ancient DNA datasets, spanning around 5,000 years, as time transects to quantify the contributions to the genome-wide variance in allele frequency change. We find that a large fraction of genome-wide change is due to gene flow. In both cases, after correcting for known major gene flow events, we do not observe a signal of genome-wide linked selection. Thus despite the known role of selection in shaping long-term polymorphism levels, and an increasing number of examples of strong selection on single loci and polygenic scores from ancient DNA, it appears to be gene flow and drift, and not selection, that are the main determinants of recent genome-wide allele frequency change. Our approach should be applicable to the growing number of contemporary and ancient temporal population genomics datasets.

**Significance statement:** The relative contribution of random genetic drift and natural selection to the change in allele frequencies through time is a long standing question in Evolutionary Biology. We show through theory and simulation how genomic time series – such as ancient DNA datasets – can be used to decompose the genome-wide contributions of selection, gene flow, and genetic drift to allele frequency change. We apply these methods to two time time series from ancient Europeans and show that gene flow accounts for most allele frequency change over the last few thousand years, with genetic drift and not selection making up much of the rest of the contribution to genome-wide evolutionary change.

## 1 Introduction

There is a long-standing debate about the role of genetic drift versus selection in evolutionary change (Buffalo, 2021; Gillespie, 1984; Jensen et al., 2019; Kern & Hahn, 2018; Kimura, 1968; Kreitman, 1996; Sella et al., 2009). While this debate has sometimes been contentious, the answers to these questions are quantitative, describing the relative contributions of basic evolutionary forces to allele frequency change and how this differs across species and different functional categories. Estimating these contributions is complicated, in part because selection can have direct and indirect effects, where the indirect effects include “linked selection”, the impact of correlated selection at nearby selected sites (Barton, 2000; Charlesworth et al., 1993; Coop, 2016; Kaplan et al., 1989; Maynard-Smith & Haigh, 1974; Sella et al., 2009). The problem is made more difficult as we often rely on a single snapshot of contemporary genomes to tease apart multiple interacting processes (gene flow, demography, hard or soft sweeps, background selection, selective interference, etc).

Genomic time series, from museum collections and ancient DNA, offer a potent reservoir of temporal genetic data to track the changes in allele frequencies, identify selected loci, and understand the impact of other evolutionary forces (e.g. Bergland et al., 2014; He et al., 2023; Le et al., 2022; Machado et al., 2021; Mathieson & Terhorst, 2022). Ancient human DNA has already revolutionized our view of human history, revealing that large-scale population movement and gene flow are pervasive, with complex patterns of allele frequency change driven by population turnover.

Unlike genetic drift, allele frequency change due to either selection or gene flow is expected to be sustained and directional. Recent investigations have highlighted the role of gene flow and population structure in confounding our interpretation of genetic signals of selection in humans (Berg et al., 2019; Petr et al., 2019; Sohail et al., 2019; Souilmi et al., 2022). Methods to look at selection at single loci and on polygenic scores in human ancient DNA now often account for the confounding effects of gene flow. These approaches have revealed persuasive signals of selection (Field et al., 2016; Irving-Pease et al., 2022; Ju & Mathieson, 2021; Mathieson & Terhorst, 2022; Mathieson et al., 2015; Souilmi et al., 2022; Wilde et al., 2014). However, these methods only capture outlier signals, and so cannot give us a full picture of how gene flow, selection, and genetic drift have driven genome-wide change. Linked selection is thought to have a critical role in shaping patterns of genetic diversity and divergence in humans on long time scales, with an autosomal reduction in diversity of upward of 20 % from background selection alone (McVicker et al., 2009; Murphy et al., 2021). Some authors have also argued for a pervasive role of selective sweeps in shaping genome-wide patterns of diversity (Enard et al., 2014; Schrider & Kern, 2017). Thus, it is an open question how much of allele frequency change genome-wide is driven by selection in humans.

We set out to decompose the total variance in allele frequency change into the contributions of linked selection, gene flow, and drift. Unlike genetic drift, ongoing selection creates covariance in allele frequency change between non-overlapping time intervals. The use of genome-wide allele frequency change covariances in time series data has recently been proposed to identify the proportion of genome-wide change due to selection in closed populations (Buffalo & Coop, 2019). In a panmictic population, a genome-wide positive covariance between a pair of time periods indicates the compounding of allele frequency change across generations, while negative covariance can potentially result from selection pressures in opposite directions. Many different modes of selection are expected to generate these covariance patterns (Buffalo & Coop, 2020; Santiago & Caballero, 1998). This approach has been applied to experimental evolution datasets (Brennan et al., 2022; Buffalo & Coop, 2020) and to natural populations where temporal sampling is available (in *Mimulus*, oaks and cod; Kelly, 2022; Reid et al., 2023; Saleh et al., 2022). These applications, along with related methods (Bertram, 2021), have revealed that a reasonable proportion of total allele frequency change, especially in artificial selection experiments, can be attributed to widespread selection beyond just a few outliers. However, applying these methods when population structure and migration are present will give biased results, as sustained gene flow across time periods can also drive temporal covariance in allele frequency change.

Here, we develop theory and simulations to show how the effect of gene flow can be accounted for using an admixture model from a known set of sources to recover the genome-wide contribution of gene flow and linked selection. We demonstrate this approach using two European human ancient DNA time series from the UK and the Bohemian region of Central Europe to quantify the contributions of linked selection and gene flow to the total variance in allele frequency change between the Neolithic and a modern or recent time point. In both cases, we find a major contribution of gene flow to allele frequency change. After correcting for known gene flow in these time transects, we do not observe any signal of genome-wide linked selection. However, we detect a weak signal of linked selection in levels of temporal allele frequency variance in regions of the genome with low recombination and high gene density.

## 2 Results

### 2.1 Model

We consider a model of data from a population sampled at multiple discrete time points (*t ∈* [0, *T*]) from an arbitrary geographic region. This population receives gene flow, modeled as single pulses of admixture, from other known source groups through time. We follow the population allele frequency over time, *p*_*t*_, and use Δ*p*_*t*_, defined as *p*_*t*+1_ − *p*_*t*_, to denote the change in allele frequency between adjacent time points. The focal population has mean ancestry fractions 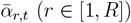 from *R* source populations that will change over time due to gene flow. We assume that allele frequencies in the source populations are constant and that good proxy samples for these sources are available, and discuss the implications of those assumptions later.

Given sample frequencies at some large set of SNPs, we can calculate the empirical variance-covariance matrix of allele frequency change for our time series, Cov(Δ*p*_*i*_, Δ*p*_*j*_), for all combinations of time intervals *i* and *j*, averaged over SNPs. In calculating these covariances we include adjustments for biases in the variance and covariance estimates due to shared sampling of an intermediate time point (see section 4.1 and appendix A). Under our model, the total variance in allele frequency change between the first sampled time point (0) and any following time point (*T*) in the time series can be decomposed into sums over time intervals of the contributions due to drift, selection and admixture:

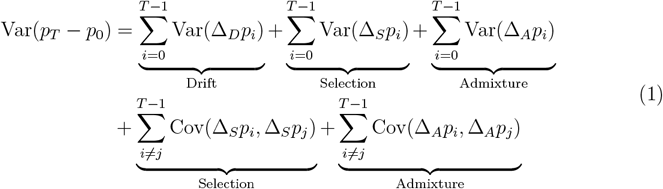

For simplicity, here we omit an interaction between drift in one time period that admixture in later time periods subsequently erases, that adds an additional term to covariances (appendix C) that we account for.

The expected variance and covariance of allele frequency change due to admixture follow from the expected allele frequency change in ancestry proportions through time. Specifically, if in the *t*^*th*^ time interval admixture changes the ancestry proportion from the *r*^*th*^ source from 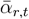 to 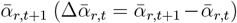, then the expected change in frequency due to admixture is 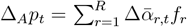, where *f*_*r*_ is the allele frequency in the source population *r*. Thus, the admixture contribution to covariance in allele frequency change between time periods *i* and *j* can be expressed as:

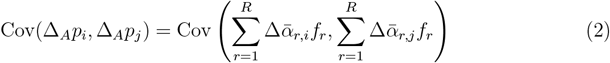

As we only have sample allele frequencies from proxies of the sources of admixture, this matrix is corrected for sampling noise biases in *f*_*r*_ (appendix B). With this admixture covariance in hand, we can now calculate the contribution of gene flow to allele frequency change, and adjust for the contribution of admixture when looking for covariances induced by selection.

We express the estimated proportion of total variance in allele frequency change attributable to gene flow (admixture) up to time *t* (*A*(*t*)) as:

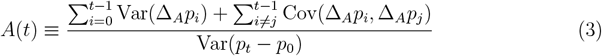

where the terms in the numerator are given by eq. (2). Note that *A*(*t*) might be an under-estimate as it excludes the contribution of gene flow from unmodeled sources, as well as gene flow events that leave the admixture proportions relatively unchanged.

The proportion of total variance in allele frequency change between 0 and *t* due to linked selection, *G*(*t*), is defined as the ratio of the total covariance due to linked selection over the total variance (Buffalo & Coop, 2019). Under our model, *G*(*t*) can be estimated by correcting the empirical covariance by the estimated covariance term due to admixture:

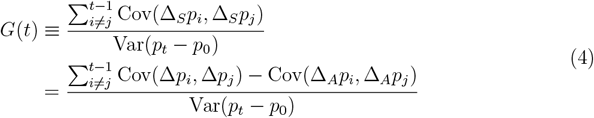

We also report an estimate of *G* not controlling for admixture (*G*_*nc*_). Note that *G* is a lower bound on the proportion as it does not account for the contribution of linked selection to the variance in allele frequency change within time periods. We attribute the residual proportion 1 − *A* − *G* of the total allele frequency variance to drift-like allele frequency change. This proportion of the temporal variance is consistent with genetic drift as it excludes covariances between time periods, which can not come from drift, and the contribution of known gene flow.

### 2.2 Simulations

To illustrate our decomposition of genome-wide allele frequency change, we simulated a simple scenario where a population receives pulses of admixture (arrows in fig. 1A and fig. S1). This repeated admixture results in positive covariances between time intervals due to the admixture-driven allele frequency change in generations 160-100 (measured before present) being in the same direction as those in generations 60-0 (fig. 1B & C). We can remove the covariance due to admixture (eq. (2)), resulting in covariances close to zero (fig. 1B & D).

**Figure 1.**
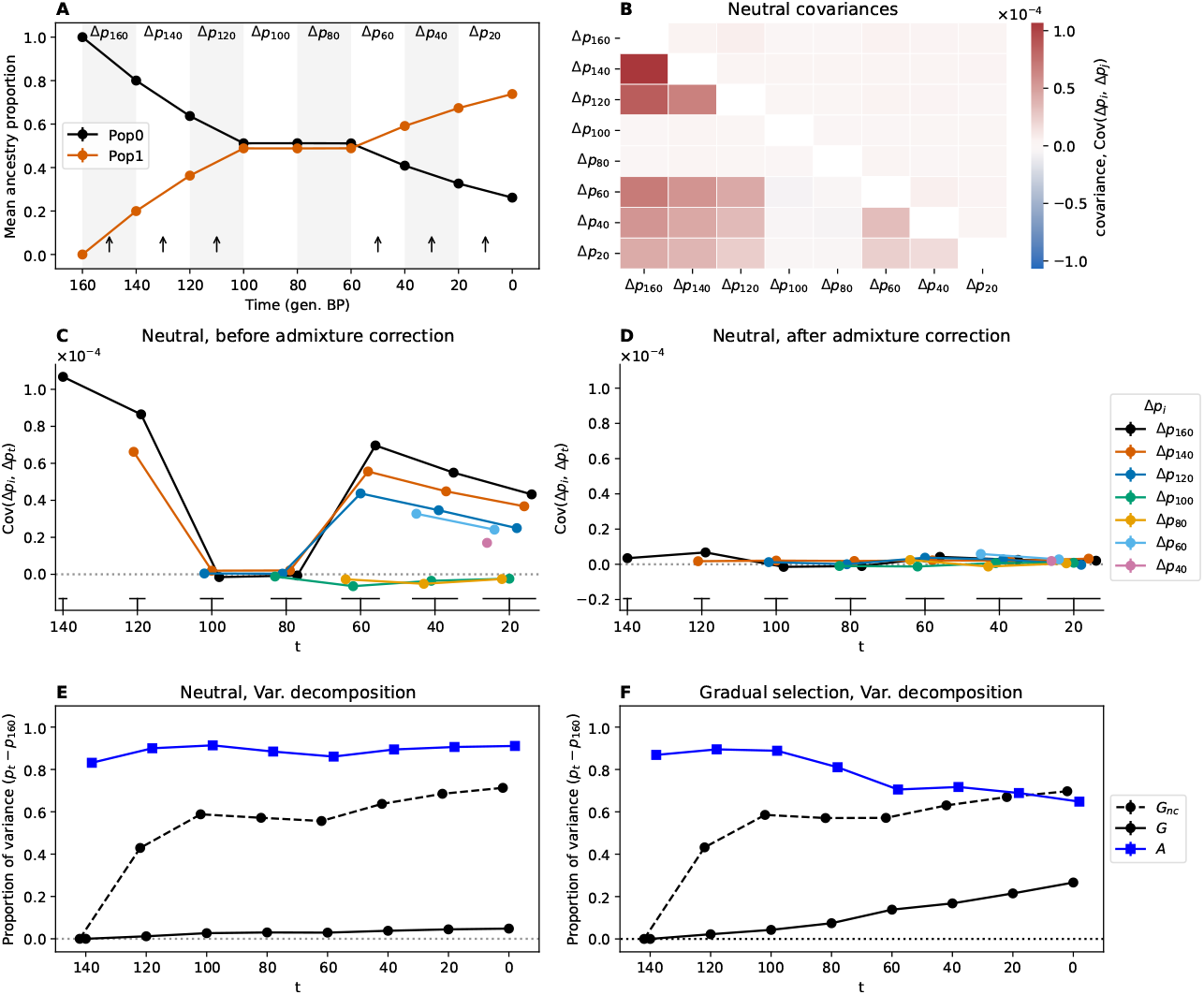
Simulation scenario of admixture between two populations (0 and 1) under neutrality (B to E) and with selection (F). **(A)** Ancestry proportions of the focal admixed population through time in generations before present. Arrows indicate the migration pulses from Pop1 (at 150, 130, 110, 50, 30 and 10 generations before present). **(B)** Covariance between time intervals. Below diagonal values are before admixture correction, above diagonal are after admixture correction. **(C)** and **(D)** pre- and post-admixture correction covariances. X-axis values are slightly shifted for visualization and the bottom lines indicate point groupings to their corresponding time. **(E)** Proportion of the total variance between the initial measured time (120) and *t* due to linked selection (*G*_*nc*_ and *G* are pre- and post-admixture correction respectively) and to gene flow (*A*). Points are slightly x-shifted for visualization. **(F)** Simulations for the decomposition of variance for neutral polymorphisms with selection around a gradually moving optimum starting at generation 140 BP for three independent traits. All points have 95 % confidence intervals (CIs) of the mean, computed using 100 replicates of the simulations (here the small CIs are hidden by their points).

We can calculate the total contribution of gene flow to allele frequency change in our simulated time series (eq. (3)) and see that gene flow accounts for much of the allele frequency change. Because the repeated gene flow creates positive covariance in allele frequency change, failing to account for this gene flow generates a spurious signal of linked selection (large non-zero *G*_*nc*_’s, dashed black line fig. 1E). However, when we account for gene flow, the signal of linked selection is almost completely removed from our neutral simulations (*G*, black line, fig. 1E). The remaining slightly non-zero *G* value in our final time period (fig. 1E) results from a slight over-correction for the covariance due to the interaction of drift and gene flow (see figs. S2 and S3F).

To illustrate the effects of selection on the covariance, we extended the above admixture simulations to include a set of loci underlying traits under stabilizing selection around a moving optimum (see section 4.2). In these simulations, we can see the proportion of neutral allele frequency change being due to selection increasing as covariances build up over time (black line fig. 1D, see also fig. S3). The effect of selection on the covariances in neutral allele frequencies is also well illustrated in a model without any gene flow (figs. S4 and S5).

### 2.3 Ancient DNA time transects

We investigated two time transects of allele frequencies in ancient humans in restricted geographical regions, the first one in the UK (Patterson et al., 2022, the England and Wales samples), and the second in the Central European region of Bohemia (Papac et al., 2021, samples from the current Czech Republic) spanning periods back to *∼* 5,600 years ago. Both these time series cover major migrations of people, where an initial mixture of early-farmer-like ancestry (EEF-like) and Western hunter-gatherer-like ancestry (WHG-like) is partially replaced by Steppe pastoralist-like ancestry (Steppe-like). This large turnover due to Steppe-like migration into Central and Western Europe was followed by a progressive increase in EEF-like ancestry over a longer time period.

We combined the Patterson et al. (2022) UK data from 793 ancient individuals with the present day GBR 1000 genomes samples (“people of European ancestry from Great Britain (GBR)”), to form a time series that runs from *∼* 5,500 years ago to the present day. Following Patterson et al. (2022) we broke the samples into 7 time periods corresponding to transitions in ancestry proportions (fig. 2C). We used the individual ancestry proportions of Patterson et al. (2022) inferred from a `qpAdm` three population model. These ancestry proportions are calculated to reflect genetic similarity to a set of samples that are pre-specified proxies for sources of ancestry. Note that an increase in a particular ancestry likely does not reflect gene flow directly from that source but rather from more nearby groups who themselves were already mixtures. In turn, each of the three major putative source ancestries was a product of admixture in the past.

**Figure 2.**
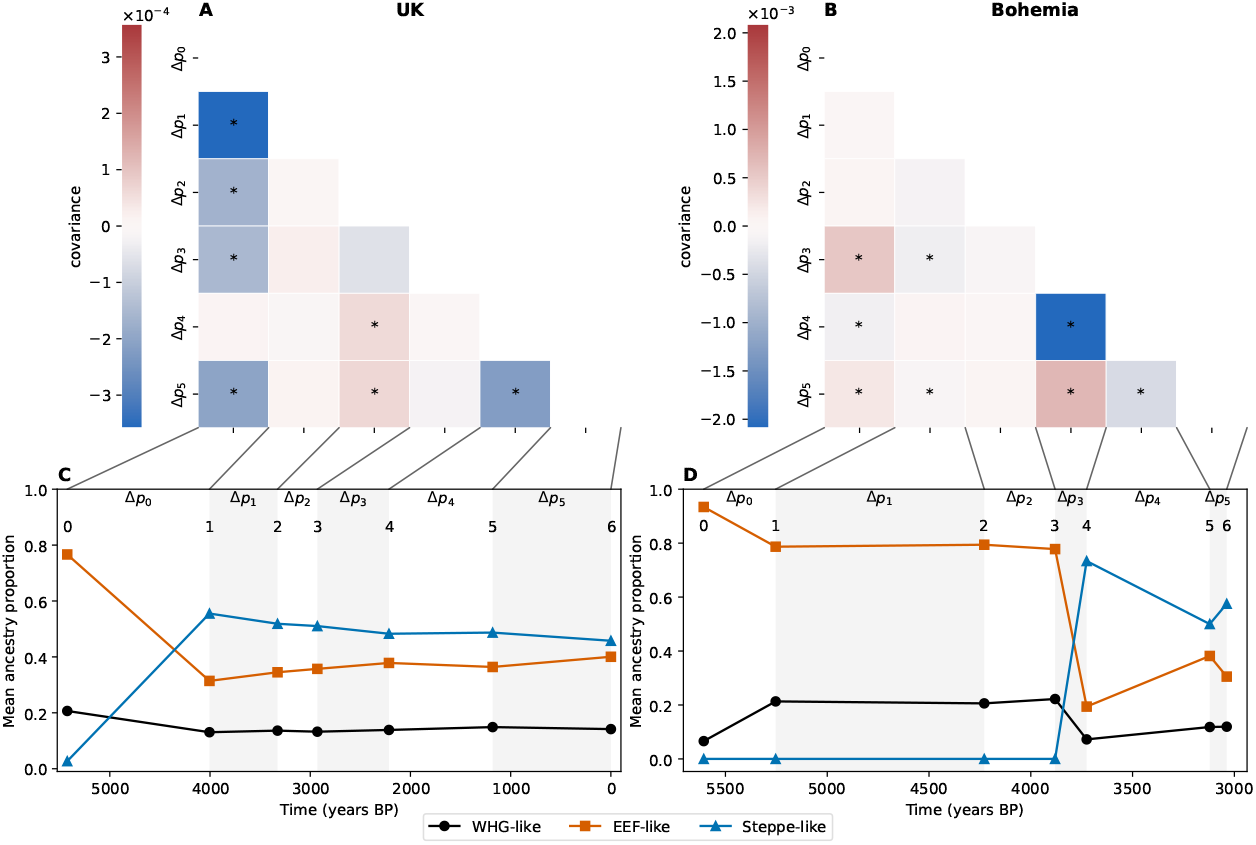
Human time series covariance matrices and ancestries (UK left column, Bohemia right column). **(A-B)** Covariance matrices with covariances significantly different from zero marked with a star. The covariances have only been corrected for sampling bias and not admixture. **(C-D)** Mean ancestry proportions from the three reference populations in the time transects samples (the mean of sample ages in each period is used for the representation).

The covariance matrix of allele frequency changes between time periods is shown in fig. 2A. The UK time transect shows several large negative covariances between allele frequency change in the first time period (Δ*p*_0_, 5424-4005 years ago) and subsequent time periods (see also black points in fig. 3A). This negative covariance largely reflects that the initial large population turnover due to Steppe-like migration (during the first time period), was followed by an increase in EEF-like ancestry (fig. 2C, Patterson et al., 2022) generating allele frequency changes in the opposite direction. After correction by admixture, covariances are strongly reduced with only a small subset differing significantly from 0 (fig. 3C). In the UK time transect, most of the total variance in allele frequency change across time is due to admixture-based changes, with 60 % of allele frequency change being driven by the Steppe-like gene flow, only for the contribution of admixture to drop gradually as the EEF-like ancestry increases due to subsequent migration(s) (fig. 3E, *G* and *A*). If we do not adjust for admixture, our estimate of the contribution of linked selection (*G*) is negative (fig. 3E, *G*_*nc*_), reflecting the negative covariances induced by admixture. After adjusting for admixture there is no signal of a long-term contribution of linked selection, with *G* not departing from zero, as there is no consistent pattern of residual positive covariances. Our empirical covariance results match those produced by a UK-like neutral simulation (Pearson and Durbin, 2023 model with UK matched admixture pulses, figs. S16 and S17). Finally, we checked for the ascertainment bias effects on our *G* and *A* estimates making use of the fact that the genotyping array consists of SNP sets discovered in different ascertainment panels. While using subsets of differently ascertained loci increases our uncertainty in our estimates, particularly for panels from more genetically distant samples, we find that our results are robust to the ascertainment scheme (fig. S8).

**Figure 3.**
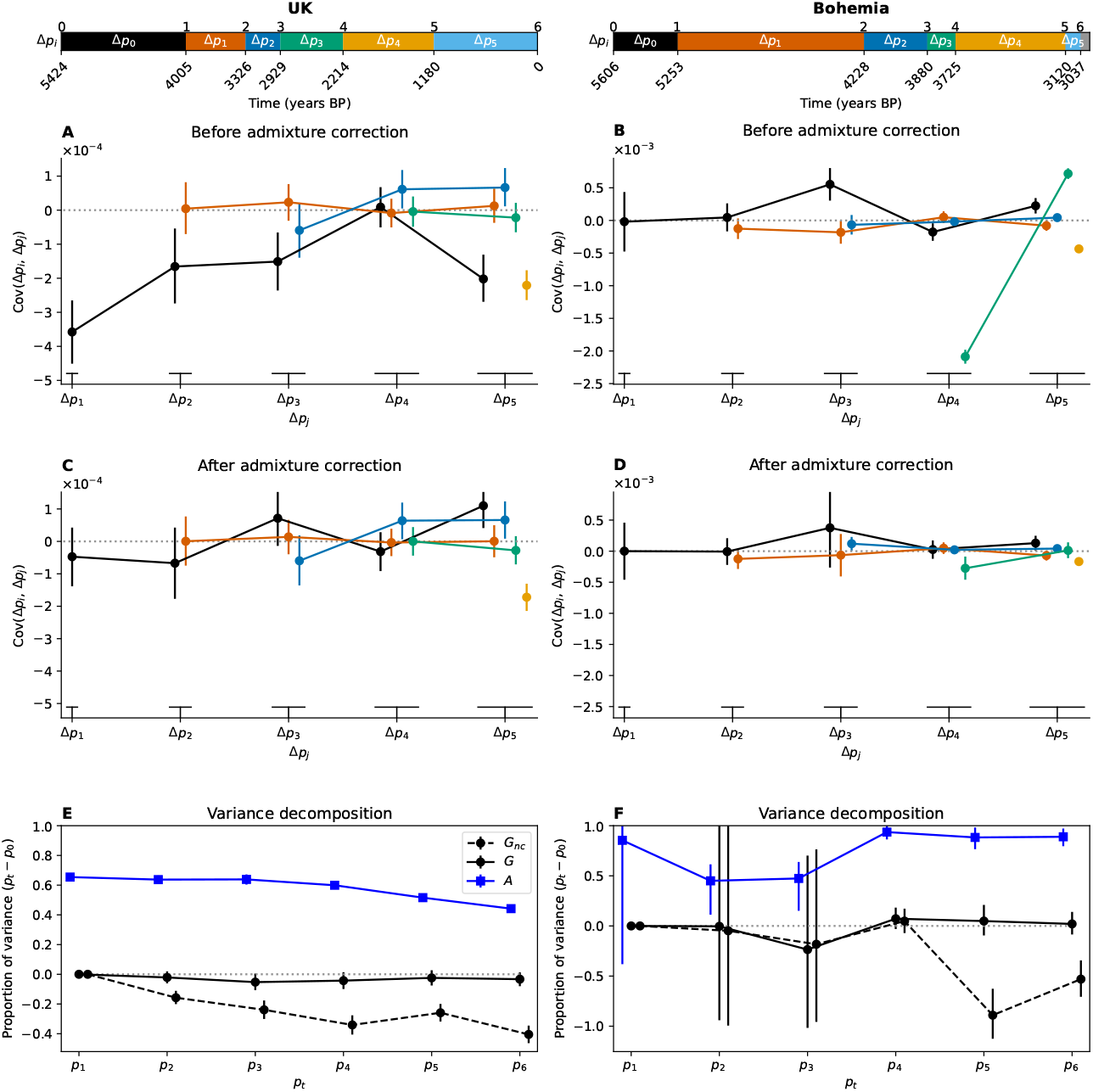
Human time series covariance corrections and variance decomposition (UK left column, Bohemia right column). The top panels show the time intervals on the years BP axis, note that the two time series have different axes. All 95 % CIs are computed through a block bootstrap procedure and represented with vertical bars. **(A-B)** Covariance values pre-admixture correction. Each line corresponds to the covariance between a first time interval (Δ*p*_*i*_, color code) and a later time interval (Δ*p*_*j*_, x-axis). **(C-D)** Covariance values post-admixture correction. **(E-F)** Proportion of the total variance, between the initial measured time (0) and time *t*, due to linked selection (*G*_*nc*_ for non-corrected and *G*) and to gene flow (*A*).

We investigated another time transect from the Bohemia region (Papac et al., 2021) spanning from 5606 to 3037 years ago, which we also split into 7 time periods following the original paper (fig. 2D). The largest covariance is negative, and an order of magnitude larger than in the UK dataset (fig. 3B), and again seems to be due to the large influx from a Steppe-like source between 3880 and 3120 years ago (between time points 3 and 4, fig. 2D) and subsequent recovery of EEF-like ancestry (between Δ*p*_3_ and Δ*p*_4_, fig. 2D). Again this large covariance due to admixture can be corrected for (fig. 3D). While the older time points have a large amount of uncertainty, due to small sample sizes (section 4.3), nearly all of the variance in allele frequency change across the full time period is accounted for by admixture (*A ∼* 0.9) and we see little evidence of allele frequency change attributable to linked selection in this Bohemian time series (fig. 3F *G*, a result that holds over SNP ascertainment scheme fig. S8).

In sum, we see little evidence, in either transect, of linked selection in the covariances in allele frequency change between time intervals, suggesting that having accounted for admixture, much of the residual change across time intervals is due to drift-like sampling processes. One caveat is that if selection operates over short time scales, e.g. selection pressures fluctuate or deleterious alleles are quickly lost, selection could generate substantial allele frequency change (variance) within time intervals but little to no covariance between the time intervals we consider.

To address the concern about the time intervals, we first reran our covariance analysis on the larger UK dataset splitting each time period in half. With this finer dissection of short-term covariances, we still see no evidence of covariance due to linked selection (fig. S12). To further explore the effect of time intervals we extended our simulations of Gaussian stabilizing selection and found that the sum of covariances decreases with the length of the time interval studied (fig. 4A), however, this effect is only pronounced when the recombination rate is low (see also fig. S6). Temporal covariances are also generated under models of background selection fig. 4B and fig. S7 Buffalo and Coop, 2020 and, while somewhat diminished, these also persist with longer sampling periods. Thus our simulations suggest that while temporal binning of ancient DNA samples will lead to lower covariances, the signal of linked selection should still be detectable.

**Figure 4.**
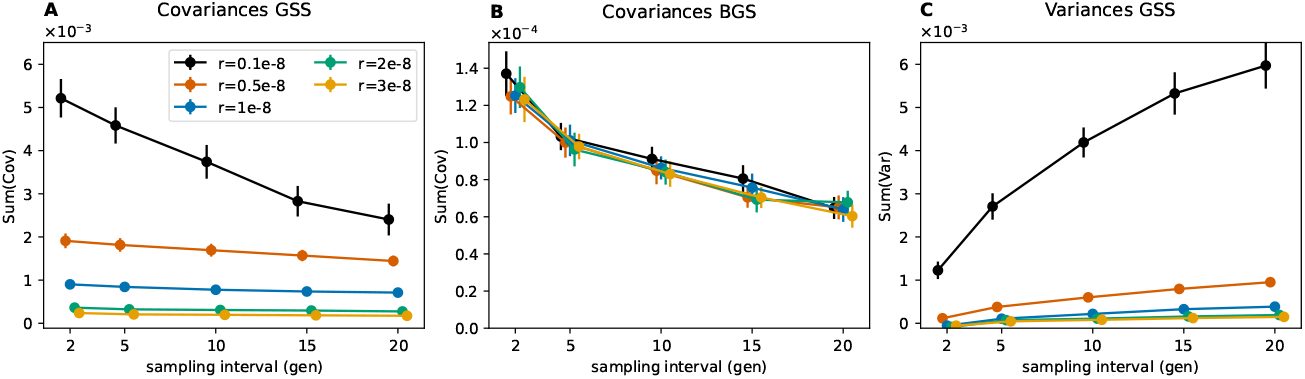
Variation of the total covariances and variances with sampling intervals (x-axis) recombination rates (color coded) for the Gaussian stabilizing selection model (GSS, **A** and **C**) and background selection (BGS, **B**). Mean and 95 % CI of the mean are plotted for either the sum of all covariances or the sum of all variances of allele frequency changes between time intervals after corrections. The sum of covariances is proportional to *G*. For each recombination rate, 100 simulations are run with sample recording at every generation starting at the creation of Pop2 in the demographic scenario with admixture (fig. S1).

One further prediction is that linked selection is expected to have larger effects in low recombination regions than high recombination regions, and in regions with a greater density of functional sites (patterns that are seen in human polymorphism datasets, Cai et al., 2009; Hernandez et al., 2011; McVicker et al., 2009). In simulations, we can see this effect, with greater temporal covariances in regions of lower recombination fig. 4A) and larger variances in allele frequency changes with lower recombination (fig. 4C). While the temporal covariances decrease with longer time interval, linked selection makes a greater contribution to the variance of allele frequency change within time intervals, so the overall signal of linked selection can be retained in the correlation of allele frequency temporal variances with recombination. To empirically examine this effect of selection we binned SNPs by their local recombination rate and a measure of the potential strength of linked selection, the B-value, which at each location in the genome combines the information of recombination rates and density of coding sites (McVicker et al., 2009; Murphy et al., 2021). In both time transects, we do not observe any significant variation in the *G* and *A* statistics recalculated in bins of recombination rate (Bhérer et al., 2017) or the B-value figs. S10 and S14). However, in the UK transect, we do see a significant increase in the total variance in allele frequency change in the lowest B-value bin (corresponding to the largest decrease in effective population size due to background selection, fig. 5A, first bin mean is above the genome-wide 95 % CI) and in the variances of change within some of the time intervals (fig. 5B). Noise in the smaller Bohemia time transect precludes seeing such effects (figs. S13 to S15). The allele frequency temporal variance increase in the UK for the lowest quintile B-value compared to the genome-wide mean is of 14.8 %, suggesting a fifth of this, *∼* 3 %, is an estimate of the genome-wide contribution of linked selection to allele frequency change.

**Figure 5.**
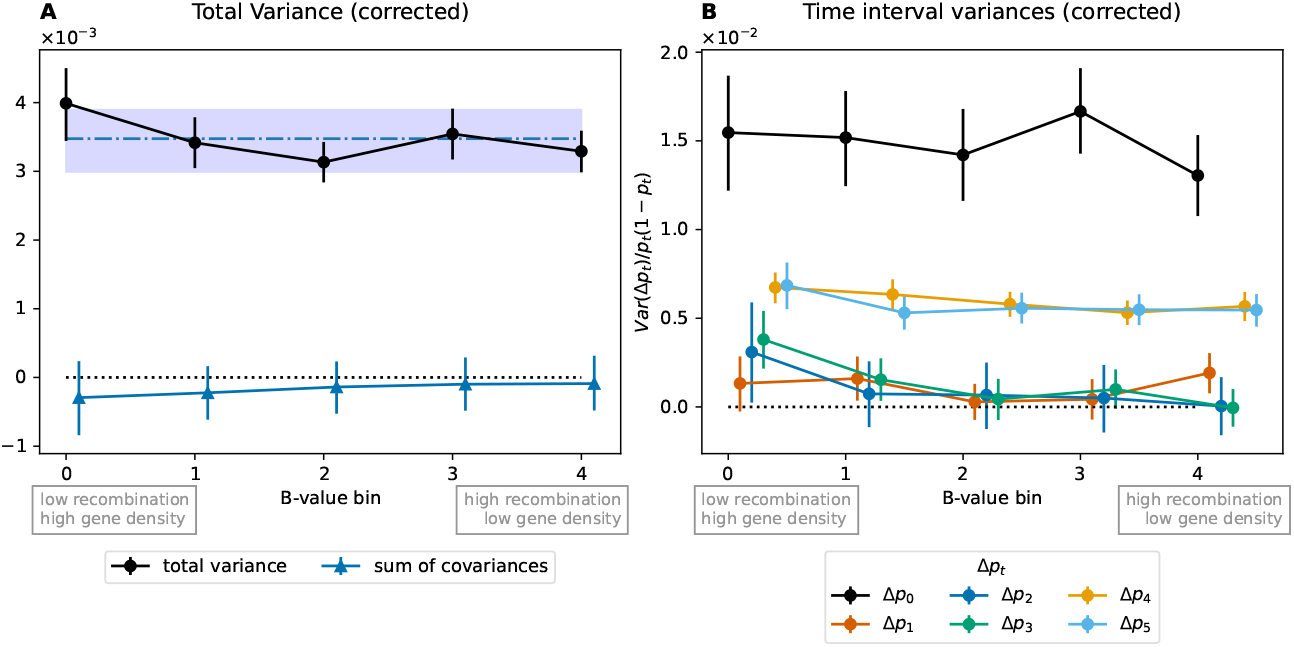
Temporal variances for the UK time series binned by a proxy for the strength of linked selection (B-value). **(A)** Total temporal variance and sum of covariance for each quintile bin of B-values. The blue dash-dotted line and interval are the genome-wide mean and 95 % block bootstrap confidence interval for total variance computed with 1/5^th^ of windows sampled on each bootstrap to be comparable to the binned values. **(B)** Variances by bin for each time interval, normalized by heterozygosity. B-value quintile bins: [0.536-0.755), [0.755-0.849), [0.849-0.902), [0.902-0.944), [0.944-0.997).

## 3 Discussion

Here we have shown how ancient DNA data can be used to decompose the contribution of gene flow, linked selection, and drift to genome-wide allele frequency change. Using two ancient DNA time transects, our results demonstrate that gene flow is the dominant force changing allele frequencies in the recent history of European human populations, and that selection-driven change is not common across the genome. This does not necessarily contradict the number of signals of temporal selection found to date, as a small fraction of loci could be subject to strong selection (e.g. Irving-Pease et al., 2022; Ju & Mathieson, 2021; Le et al., 2022; Mathieson & Terhorst, 2022; Mathieson et al., 2015; Wilde et al., 2014). Indeed, some of these methods apply similar admixture adjustments as ours, but look for genome-wide outliers and so only detect strong selection on single loci (e.g. Mathieson and Terhorst (2022) expect to be able to detect selection coefficients *>* 0.02). Another set of approaches looks for ancient selection on polygenic scores constructed from genome-wide association studies (Le et al., 2022; Mathieson & Terhorst, 2022). These approaches account for admixture and can detect subtle shifts at loci in ancient DNA, but rely on the fraction of genetic variation for specific traits captured in modern-day samples (Yair & Coop, 2022). Thus, our genome-wide method is complementary to both time series outlier approaches as well as phenotype-motivated approaches.

The large contribution of gene flow to evolutionary change in the past few thousand years is not surprising given the dynamic picture of population movement that has emerged from ancient DNA. Our results provide additional evidence that allele frequency changes are well fit by relatively simple admixture models, and strengthens the view that multiple migrations events throughout the history of European Human populations have played a preponderant role in the composition of modern populations. The lack of a substantial contribution of linked selection is perhaps more surprising. Linked selection has been estimated to account for upward of 20 % reduction in long-term patterns of human diversity (under models of background selection, McVicker et al., 2009; Murphy et al., 2021), and so we should expect a similar portion of the variance in allele frequencies to come from linked selection. Much of this effect should manifest itself in the compounding of positive covariances between allele frequency changes across the generations. While this effect has been seen empirically in selection experiments and in some natural populations, we currently do not see any evidence of this in humans. One possibility is that the time periods we consider are not long enough for strong covariances to build up, as the long-term patterns of linked selection reflect dynamics over coalescent time scales of hundreds of thousands of years. In contrast, the other possibility is that negative selection generating background selection is fast enough that it does not contribute to covariances among the time periods used here. However, under this latter interpretation, we should see higher allele frequency variances in regions predisposed to stronger linked selection, but we see this effect only weakly when partitioning loci by B-value. Larger collections of ancient DNA will allow better temporal resolution of allele frequency covariances, which could be combined with more individual-level approaches to avoid the need for sample lumping in time periods. It is also possible that some signals of linked selection may be washed out at the fine geographic scale of our time series, as our time series approach may partially be picking up ephemeral change which may average out over the much larger meta-population within which our time series are embedded.

Our approach uses ancestry proportions from ancient DNA for the three major inferred waves of gene flow into Europe. The sparsity of ancient DNA means that we rely on the ancestry proportions of relatively small samples of ancient individuals to be representative of people living in the past. However, the periods that we divide our samples into reflect reasonably well-established periods in the peopling of the regions. We also rely on allele frequencies in a set of samples as proxies for sources of gene flow. As we discuss below, the misspecification of the sources of gene flow may appear as evolutionary change within our focal time series. One future extension might be to use principal components analysis to learn about major axes of population structure involving samples in a time series and then to regress these PCAs out of our genotypes to account for variation in ancestry composition in a more model-free manner.

Our admixture correction seems to perform well on time intervals involving the large ancestry shift in Steppe-like and EEF-like ancestry (compare black points between fig. 3A and C). However, we see several negative covariances that remain after adjustment for admixture (fig. 3A and C). In principle, these could reflect fluctuating selection, but that seems unlikely given the general lack of other evidence of selection. Rather, these covariances could reflect that our proxies for gene flow sources only capture part of the allele frequency change driven by gene flow. Indeed, the increase of EEF-like ancestry in the UK population is driven by subsequent migration(s) from populations similar to the UK but with a higher proportion of EEF-like ancestry, probably from mainland Europe. Therefore, modeling the increase of EEF-derived alleles with the ancestral EEF allele frequencies might not fully account for the impact of migration. More detailed modeling with admixture graphs and tree sequences could help better resolve the sources of gene flow in time series (e.g. Allentoft et al., 2022; Irving-Pease et al., 2022; Pearson and Durbin, 2023, although such inferences may currently not be fully robust, Maier et al., 2023).

We attribute residual variance after accounting for gene flow and temporal covariances to drift-like processes. Genetic drift from the compounded sampling of parents to form each generation in our geographic area will obviously contribute to this. However, as our focal geographic areas are not homogeneous populations, small changes in ancestry composition over time not captured by larger-scale admixture analyses might be captured as drift-like processes. Finally, as we take a fixed sample to reflect the allele frequencies in the sources of gene flow, change in the actual groups contributing to gene flow can also contribute to the signal of drift; e.g. if the allele frequencies in the source of EEF-like ancestry early differs from those contributing EEF-like ancestry later in the time series, that will appear as drift-like change in our focal time series. Our drift-like change in allele frequency is small, corresponding to relatively large estimates of temporal effective population sizes (see section 4.3). However, further work is needed to separate the long-term effect of drift and the combined contribution of other drift-like processes to our estimates of allele frequency change.

Extensions of our approach to larger geographical areas would allow the contributions of local genetic drift and migration among regions to be more fully explored. Such analyses would also pose an interesting set of modeling challenges to measure evolutionary change across spatially spread populations experiencing both local migration and more long-range gene flow events.

Finally, while a large body of ancient DNA work has focused on humans, ancient DNA and museum datasets for a wide range of other organisms are also being generated (e.g. dogs, Bergström et al., 2020; horses, Orlando, 2020; sticklebacks, Kirch et al., 2021; chipmunks, Bi et al., 2019; *Amaranthus*, Kreiner et al., 2022). The spread of ancient DNA and museum DNA research as well as more widespread usage of genome-wide sequencing to temporally monitor contemporary natural populations will generate a rich set of resources of time series data. This offers the chance for comparative studies to decompose the contribution of different forces to genome-wide evolutionary change across systems, time scales, and ecological and selection regimes.

## 4 Materials and Methods

### 4.1 Calculating the covariance matrix

We bin our samples into a set of discrete time points and then calculate the allele frequency change at SNP *l* between adjacent time points, *t* and *t* + 1, Δ*p*_*t,l*_. We then calculate the empirical variance-covariance matrix of these allele frequency changes for all time points averaged across SNPs. We denote the raw covariance matrix by **R**. We wish to quantify the expected contribution of admixture to this matrix, but in doing so we also have to correct for sampling noise in both the time series allele frequencies and sources of admixture. The corrected covariance matrix is given by

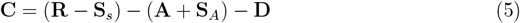

**S**_*s*_ is the expected matrix of biases from using sample frequencies in calculating the empirical covariance matrix (see eq. (6) below). Our admixture adjustment, **A**, is the expected admixture variance-covariance matrix (see eq. (2)), where proxy samples are used as references for the admixture sources. The matrix **S**_*A*_ is the expected bias in the admixture matrix due to the sample noise from using sample frequencies in our admixture correlation (see eq. (7) below). Finally, **D** is the expected drift/admixture interaction matrix (see appendix C).

Here we calculate the sampling biases in the specific case of pseudohaploid data in line with the ancient DNA datasets considered in this paper (appendix A), using *n*_*i*_ for the haploid sample size at time point *i* and 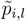 the sample frequency at SNP *l*. The sampling noise from taking a small sample of individuals inflates the variance of allele frequency change and shared sample between adjacent time points creates covariance

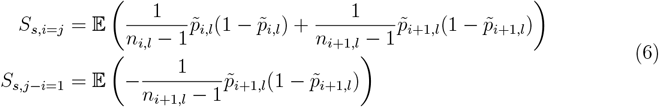

with all other terms in the matrix set to zero (appendix A and Buffalo and Coop, 2019). These matrices are calculated as an average over all our SNPs. Second, sampling noise is also present in frequencies of the samples used as proxies for admixture, and so this biases the admixture expectation as the same reference samples are used for multiple time points (appendix B):

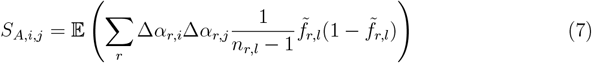

following eq. (B.3) with *α*_*r,i*_ the admixture proportion from reference population *r* at time *i*, and 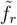 the empirical allele frequency in the reference population *r*.

Finally, the simple admixture covariance expectation is missing a term due to shared drift variances between time intervals. This can be estimated as shown in appendix C and requires the assumption that only one parental population is contributing to gene flow during each time interval. **D** is given by eq. (C.9) and is dependent on the estimated drift variance terms in parental populations and admixture proportions at each time step common between two time points.

### 4.2 Simulations

We used the Demes format to write inter-operable demographic scenarios (Gower et al., 2022). This allowed us to run the same model with either `msprime` for neutral simulations (Baumdicker et al., 2022) or `SLiM` (v3.7) for simulations including selection (Haller & Messer, 2019). Results were recorded as treesequences and analyzed in `Python` using `tskit` (Kelleher et al., 2018). All results are based on 100 replicates of each scenario. The simulations pipeline was built with `snakemake` (Mölder et al., 2021) and can be found in the zenodo archive https://doi.org/10.5281/zenodo.8093105 and includes the version of all software used.

In the main text simple scenario, an ancestral population splits into the source populations 1,500 generations before present (BP). All populations are kept at a constant size of 10,000 diploid individuals. 200 generations BP our focal population that receives the admixture pulses is created from the first parental population (pop0). Pulses of admixture from pop1 happen at regular 20 generations interval starting at 150 and finishing at 10 generations BP. We sample 30 individuals 10 generations before and after pulses in our focal population. For our admixture sources, we sample 30 individuals from each parental population at 200 generations BP for allele frequency computations. Samples are rendered pseudohaploid to mimic ancient DNA results (though no missing data was inserted). A census event of all populations is performed in the source populations when the admixed population is created to allow us to compute the admixture proportions of all descendants.

We simulated a chromosome 100 Mbp in length with a mutation rate of 1 *×* 10^−8^ per bp and per generation and a uniform recombination rate of 2 *×* 10^−8^.

To simulate linked-selection in `SLiM` (Haller & Messer, 2019), we considered three independent polygenic traits with alleles having a random effect size of *±*0.01 evolving under a model of stabilizing selection around an initial optimum of 0 for each trait. The fitness landscape is a Gaussian function centered on the optimum with a variance, *V*_*s*_, of 1. The optimum is gradually shifted from 0 to 3 standard deviations between 140 generations BP to the present similarly in all extant populations (by steps of shift/time). The ancestral population has a burn-in of 0.1N generations in `SLiM` and the complete ancestral history has been recapitated with `pyslim` (Ralph et al., 2023). Mutations under selection are not used in the downstream analyses and neutral mutations are added a posteriori with `msprime`. We note that these simulations are not intended to mimic a particular selection scenario, as the density of loci underlying different traits per chromosome is unknown. Rather the parameters were chosen to generate results where both admixture and selection made comparable contributions for illustration purposes.

Finally, to investigate the role of sampling intervals and recombination rates we repeated the above Gaussian stabilizing selection simulations with varying recombination rates ([0.1 *×* 10^8^, 0.5 *×* 10^8^, 1 *×* 10^8^, 2 *×* 10^8^, 3 *×* 10^8^]) approximately spanning the range of human recombination rates, and analyzing the outputted treesequences at different time sampling intervals ([2, 5, 10, 15, 20]). We also simulated background selection using a model similar to Buffalo and Coop (2020) in `SLiM` where we use several deleterious mutations per haploid genome per generation *U* = 1 and a negative selection coefficient *s* = − 0.1. The rest of the model and analysis pipeline is similar to the Gaussian stabilizing selection case above.

### 4.3 Ancient DNA analyses

We used two datasets from Patterson et al. (2022) and Papac et al. (2021) for ancient DNA time transects in the UK and the Bohemian region respectively. Data from those papers were downloaded from the indicated sources and merged with a set of parental population proxies and modern samples from the AADR v50.0 1240k dataset (Mallick et al., 2023 using data from Allentoft et al., 2015; Bergström et al., 2020; Brace et al., 2019; Consortium, 2015; Coutinho et al., 2020; Fu et al., 2016; Haak et al., 2015; Jones et al., 2015; Lamnidis et al., 2018; Lazaridis et al., 2016, 2017; Lipson et al., 2017; Mallick et al., 2016; Margaryan et al., 2020; Martiniano et al., 2016; Mathieson et al., 2015, 2018; Mittnik et al., 2018; Narasimhan et al., 2019; Olalde et al., 2018; Patterson et al., 2012; Raghavan et al., 2014; Rivollat et al., 2020; Scheib et al., 2019; Schroeder et al., 2019; Skoglund et al., 2015; Villalba-Mouco et al., 2019; Wang et al., 2019). Modern samples were used to provide a modern time point in the UK time transect. The data analysis `snakemake` pipeline can be found in the zenodo archive https://doi.org/10.5281/zenodo.8093107 and includes the version of all software used.

Individuals from each time period defined in the original analyses are pooled together to compute allele frequencies and the mean estimated age is taken as the time point date. For the UK dataset (Patterson et al., 2022), we merged the published data with AADR v50.0 (providing modern samples and parental population proxies), and with data from Fowler et al. (2022) to access 10 individuals missing from the other datasets. Only loci with more than 10 genotypes in each time point grouping and more than 5 genotypes in reference populations were kept, resulting in 474,554 SNPs kept over the initial 1,135,618. We used combined filters 0 and 1 from Table S5 of Patterson et al. (2022) as our quality and relevance filtering. This resulted in sample sizes of [37, 69, 26, 23, 273, 38, 62] for periods labeled [‘Neolithic’, ‘Chalcolithic/EBA’, ‘Middle Bronze Age’, ‘Late Bronze Age’, ‘Iron Age’, ‘Post Iron Age’, ‘Modern’] and mean non-missing genotypes across all SNPs of [25.5, 46.3, 19.6, 14.8, 208.2, 25.5, 59.9]. Mean sample dates for those periods are [5424, 4005, 3326, 2929, 2215, 1180, 0] years BP. Reference sample sizes are [18, 21, 18] for groups labeled [‘WHGA’, ‘Balkan_N’, ‘OldSteppe’] in the dataset interpreted as WHG-like, EEF-like, and Steppe-like. Those reference samples have mean non-missing genotypes across all SNPs of [7.4, 15.4, 12.4] respectively.

For the Papac et al. (2021) dataset, only loci with more than 2 genotypes in each time point grouping and more than 2 genotypes in reference samples were kept, resulting in 461,844 SNPs kept over the initial 1,150,639. Sample sizes in this dataset are [3, 5, 29, 14, 48, 59, 84] for periods labeled [‘Neotithic’, ‘Proto-Eneolithic’, ‘Early Eneolithic’, ‘Middle Eneolithic’, ‘Corded Ware’, ‘Bell Beaker’, ‘Unetice’] and mean non-missing genotypes across all SNPs of [3., 3.7, 23.4, 10.9, 33.1, 41.9, 55.1]. Mean sample dates for those periods are [5607, 5253, 4229, 3880, 3726, 3120, 3037] years BP. Reference sample sizes are [4, 17, 15] for groups labeled [‘WHG’, ‘Anatolia_Neolithic’, ‘Yamnaya’] in the dataset interpreted as WHG-like, EEF-like, and Steppe-like. Those reference samples have mean non-missing genotypes across all SNPs of [3.5, 13.1, 9.0] respectively.

We used admixture measures from both published papers produced by the `qpAdm` method, extracted from the supplementary Table S5 from Patterson et al. (2022) and Table S9 from Papac et al. (2021). In concordance with the literature on European Human demographic history during the last 5000 years, we consider the simplest three-way admixture between populations genetically most similar to European early farmers (EEF-like, early migrants from Anatolia), Western hunter-gatherers (WHG-like) and individuals associated to the Steppe pastoralists Yamnaya culture (Steppe-like).

We computed confidence intervals around estimates by block bootstrap sampling of windows of 1000 SNPs along the genome. Statistics computed for each window were re-sampled 10^4^ times with replacement and a 95 % confidence interval was computed by the pivot method as in Buffalo and Coop (2020). Statistics were computed through a weighted average to account for variability in the number of SNPs in each window (windows at the end of chromosomes often do not contain the required number of SNPs). When dealing with ratio statistics (like *G*), we computed separately the numerators and denominators and used the ratio of the weighted averages for the final values.

Each dataset was transformed from the `eigenstrat` to the `sgkit` format through a `plink` (Chang et al., 2015) conversion step. Sex chromosomes were removed from the datasets. To investigate the correlation of our statistics with recombination or background selection, we incorporated in the dataset recombination rates (Bhérer et al., 2017, sex-averaged version) and B-values (Murphy et al., 2021) for each SNP – by using the value of the window the SNP was in. We split all SNPs into five quantile bins and computed *G* and *A* proportions for each one, as well as the variance and covariances.

We can compute a simple estimate of the diploid effective population size, 2*N*, by equating the expected variance due to drift after *t* generations,

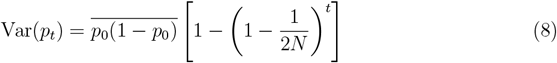

with the residual variances for each time period in the studied datasets (having adjusted for the variance due to admixture and sampling). Using a generation time of 30 years, and using the number of generations between the mid-points of each time interval for the UK dataset we obtain 2*N* = [351472, 3204693, 2827982, 2041971, 361192, 203114] for each time interval. For the Bohemia dataset time intervals, we get 2*N* = [ 768209, 2576065, 1392381, 1181146, 1275489, 1167632]. We note that these effective population size estimates are only approximate as they do not account for the more continuous distribution of sampling times present in the data.

## Data availability

No new data was produced for this work. Analyses pipelines are available at doi: 10.5281/zenodo.8093105 for simulations and at doi: 10.5281/zenodo.8093107 for the ancient DNA data. Those analyses rely on a custom helper python module available at doi: 10.5281/zenodo.8093101.

## Acknowledgments

We thank members of the Coop lab, Vince Buffalo, and Joshua Schraiber for helpful discussions. We also thank the editor and reviewers for helpful comments during the review process. AS and GC were supported by the National Institute of General Medical Sciences of the National Institutes of Health (NIH R35 GM136290 to GC).

## Appendix

### A Pseudohaploid sampling

Following Buffalo and Coop (2020), the observed variance in allele frequency at time *t* can be decomposed with the law of total variance:

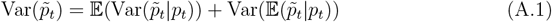

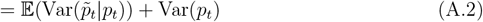

This gives us a way to correct the observed variance for sampling noise.

Pseudohaploid representation is common in ancient DNA data to avoid errors when calling heterozygotes. Most often, one read (and therefore allele) is selected randomly among the mapped reads for each individual at a given position. Pseudohaploid calling can be modeled as a binomial sampling. We consider sampling *n*_*t*_ individuals in a population with frequency of the alternate allele *p*_*t*_ at time *t*. Pseudohaploid calling is equivalent to each individual drawing one allele from the pool of alleles. We define *X*_*t*_ *∼* Binom(*n*_*t*_, *p*_*t*_) the number of alternate alleles sampled and 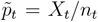. Then the sampling noise is

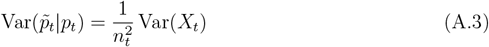

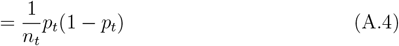

Correction of the variance 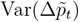 is carried out by subtracting both sampling variances 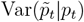 and 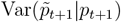. As in Buffalo and Coop (2020), the covariances between two overlapping time intervals, 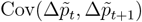 are negatively biased by the shared sampling noise in *p*_*t*+1_, and this needs to be corrected by adding the shared time point sampling variance 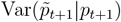 back in. For these corrections, we need an unbiased estimator of the half heterozygosity. We define the sample heterozygosity as 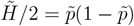, then

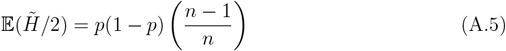

Therefore

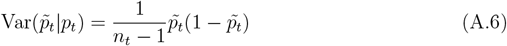

Similarly, if needed, we can compute the diploid sampling bias estimator:

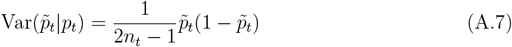

### B Pseudohaploid sampling noise in reference populations

Let’s consider the allele frequency of reference population *r*

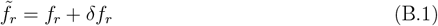

The observed allele frequency is equal to the true allele frequency (*f*_*r*_) plus sampling noise.

Therefore

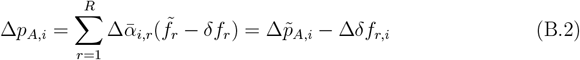

Decomposing Cov(Δ*p*_*A,i*_, Δ*p*_*A,j*_) as 𝔼 (Δ*p*_*A,i*_Δ*p*_*A,j*_) − 𝔼 (Δ*p*_*A,i*_) 𝔼 (Δ*p*_*A,j*_) and remembering that 𝔼 (Δ*δf*_*r,i*_) = 0, we end up with 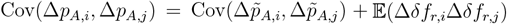

Therefore

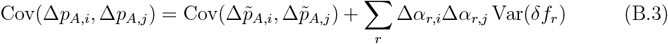

with Var(*δf*_*r*_), the variance of sampling noise in the pseudohaploid case, equal to 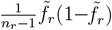 (similar to appendix A).

### c Accounting for drift in the admixture correction

We consider a simple model where only the focal admixed population experiences drift and parental populations from which gene flow occurs are not. This is in line with our use of a single proxy sample for the sources of admixture. Under this model drift happening in any of our contributing populations is absorbed into the drift observed in the focal population. Drift that occurs in one time interval can partially be erased by admixture in subsequent time intervals. This interaction between drift and gene flow generates additional covariance that needs to be accounted for.

Let *f*_*r*_ be the frequencies of *R* parental populations for a particular SNP. At time 0, an admixed population of frequency *p*_0_ is established with ancestry proportions *q*_*k*0_ from any of the *R* populations. Subsequently, between time points *t* and *t* + 1, this admixed population can receive a migration pulse from any of the *R* populations, where a proportion *α*_*r,t*_ of individuals in the focal population are replaced by migrants. Between time intervals *t* and *t* + 1, drift happens changing the frequency by Δ*d*_*t*_. *p*_*t*_ is the allele frequency at time *t* of our admixed population. *q*_*r,t*_ are the ancestry proportions at time *t* of this admixed population.

We define the proportion of individuals replaced by admixture as:

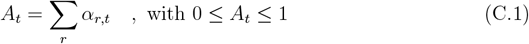

The change in allele frequency can then be written as:

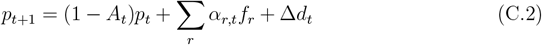

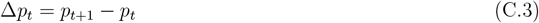

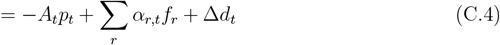

We can expand this out in terms of the change in allele frequency due to admixture and drift in the preceding time periods:

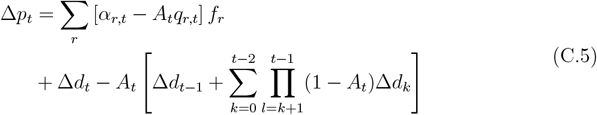

We can express the ancestry fraction from source *r* at time *t* in terms of the change due to admixture in previous time periods:

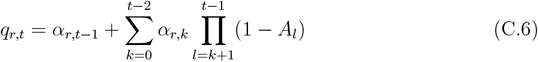

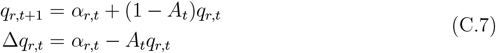

As a constraint, if at each time step there is only one *α*_*rt*_ *>* 0, then it simplifies to:

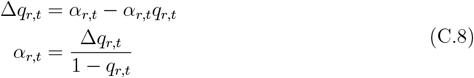

allowing us to compute the admixture fraction from the ancestry proportions.

When computing the covariance between two time intervals *i* and *j* (*i < j*), a composite drift term depending on the admixture pulses will be shared between all time intervals between times 0 to *i* between the two Δ*p*_*i*_ and Δ*p*_*j*_. This expected admixture-drift term *D* can be computed as:

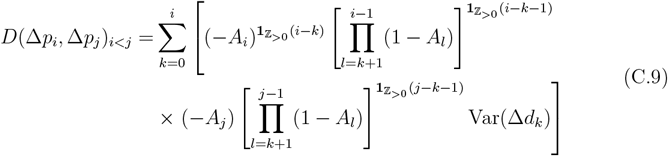

with

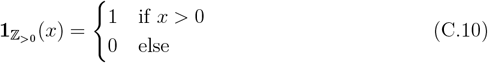

*D* can then be subtracted from the empirical covariance to remove this effect.

To compute *D*, individual time interval drift variances need to be estimated (Var(Δ*d*_*k*_)). We can use the fact that the variances at each time interval can be decomposed as a linear combination of drift variances to solve for them. Solving is only possible when considering that only one parental population is contributing to gene flow at a given time to be able to estimate the values of individual *α*_*r*_ (eq. (C.8)). The system for 0 *≤ i ≤ t* is of the form:

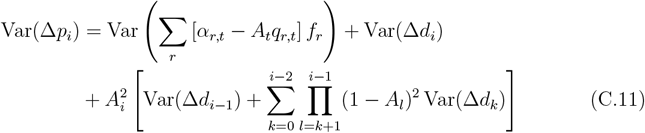

