## Supplementary_information for "The contribution of gene flow, selection, and genetic drift to five thousand years of human allele frequency change"

### Supplementary Information for Simon & Coop 2024, doi: https://doi.org/10.1101/2023.07.11.548607

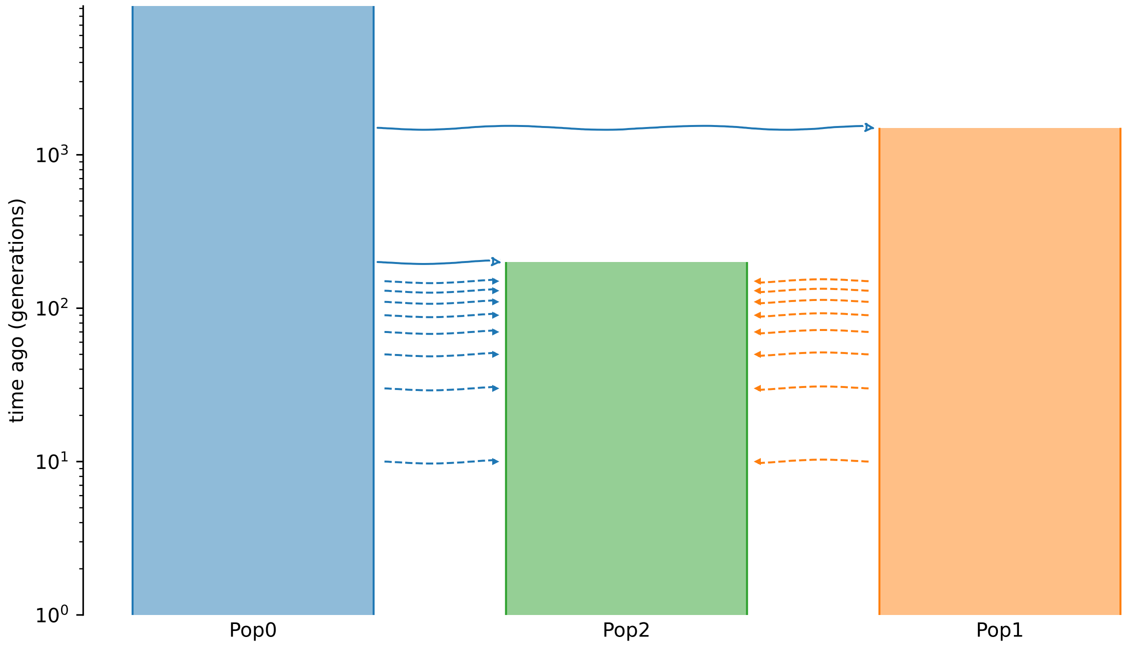


Figure S1: Demographic scenario with two ancestral populations of 10,000 individuals and an admixed population of the same size receiving gene flow. Some arrows represent null pulses, but see fig. 1A.

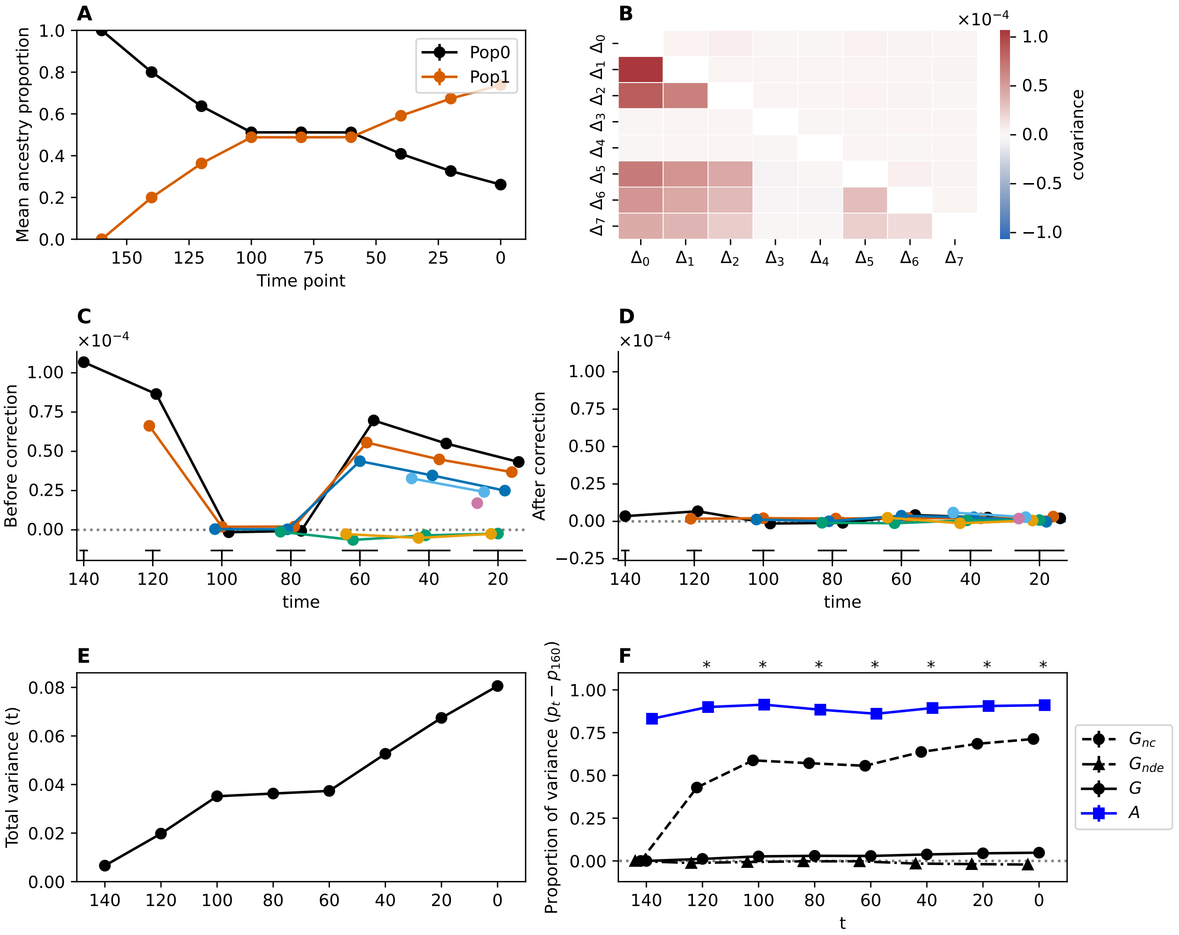


Figure S2: Neutral simulation. Similar figure to fig. 1 with total variance **(E)** and G without drift error correction, $G_{nde}$, plotted **(F)**. Stars in **(F)** indicate the absence of overlap with zero of the 95 % CI of the mean for $G$.

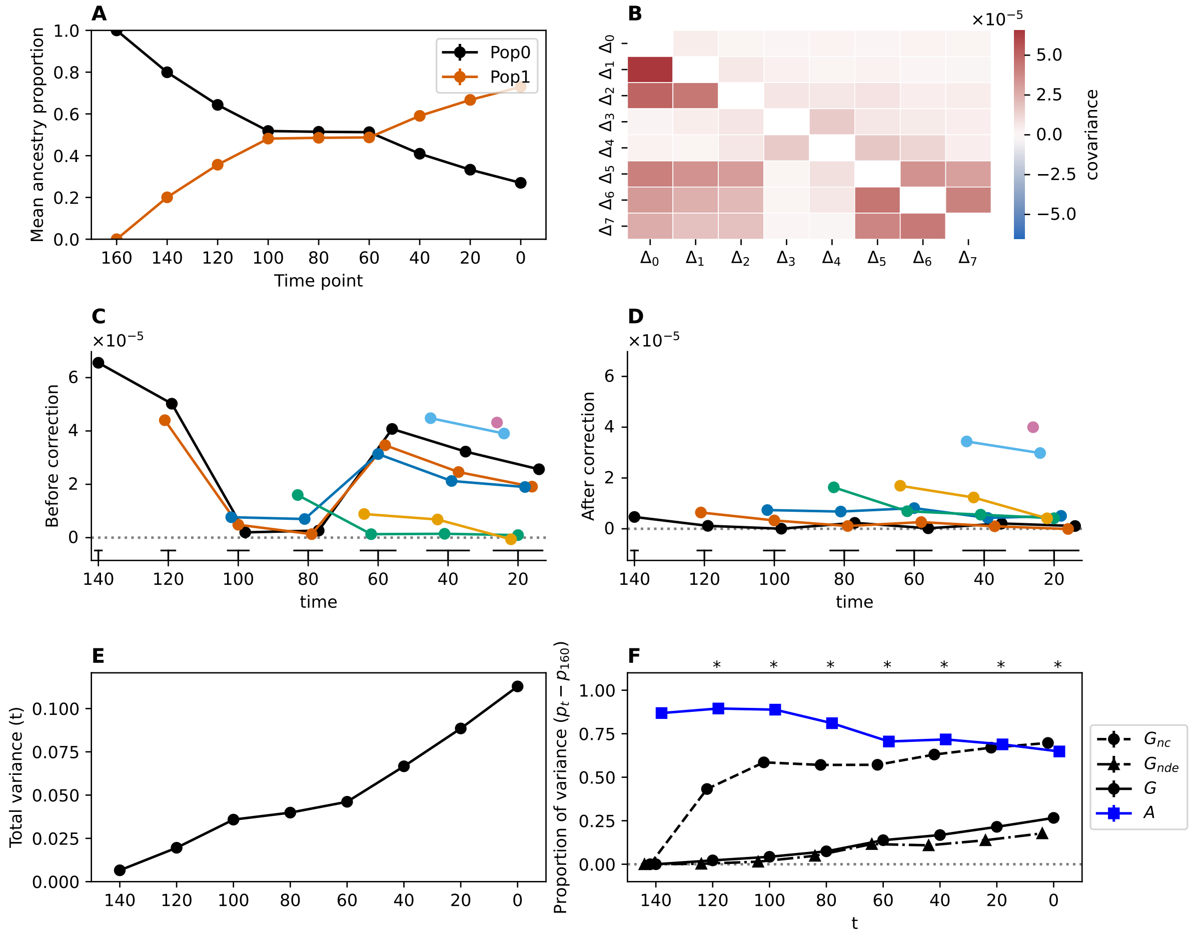


Figure S3: Same as fig. S2 with selection. Stabilizing selection around an optimum for three traits, gradually moving to a value of 3 between generation 140 and 0 BP. See simulation methods for further details. This figure is the complete presentation for the data shown in fig. 1F. Stars in **(F)** indicate the absence of overlap with zero of the 95 % CI of the mean for $G$.

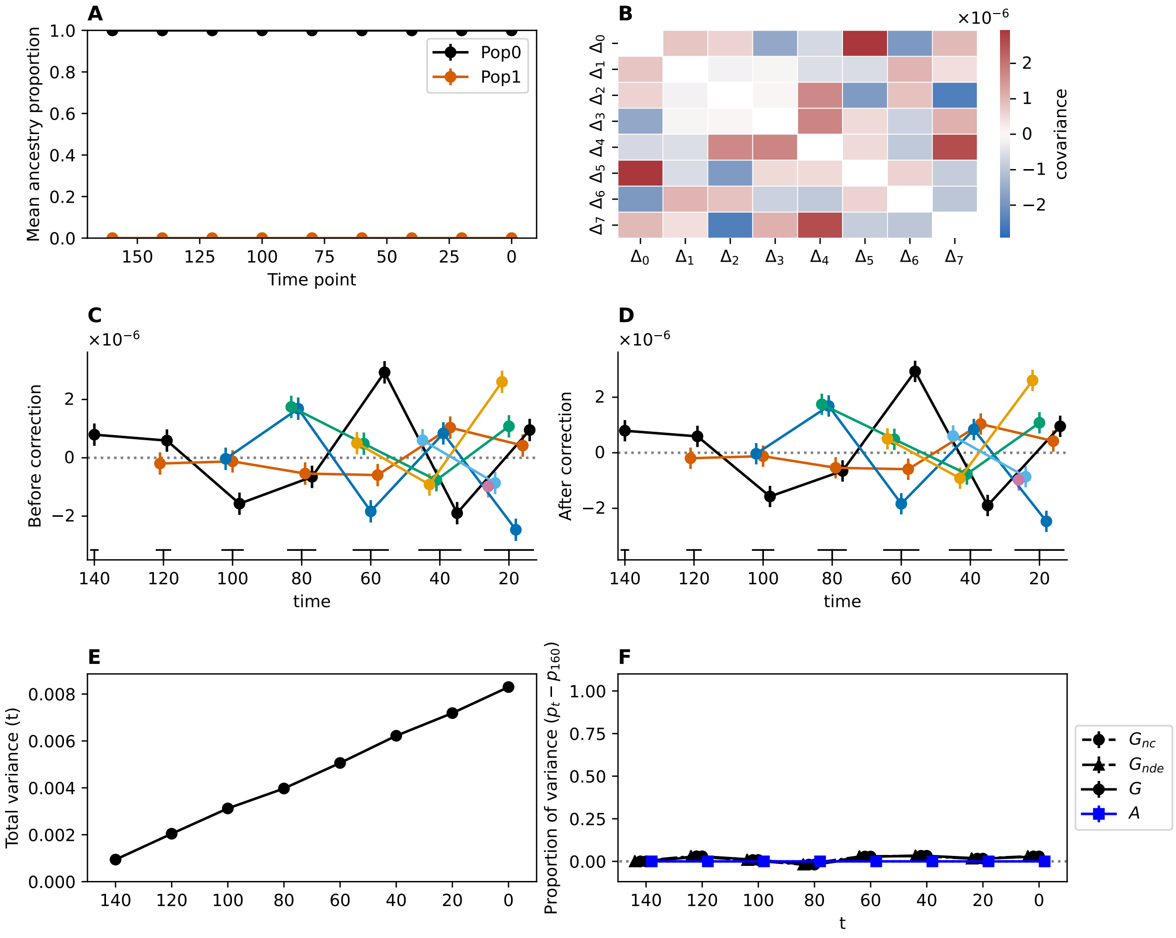


Figure S4: Demographic scenario with no gene flow or selection (all fig. S1 pulses null), neutral mutations.

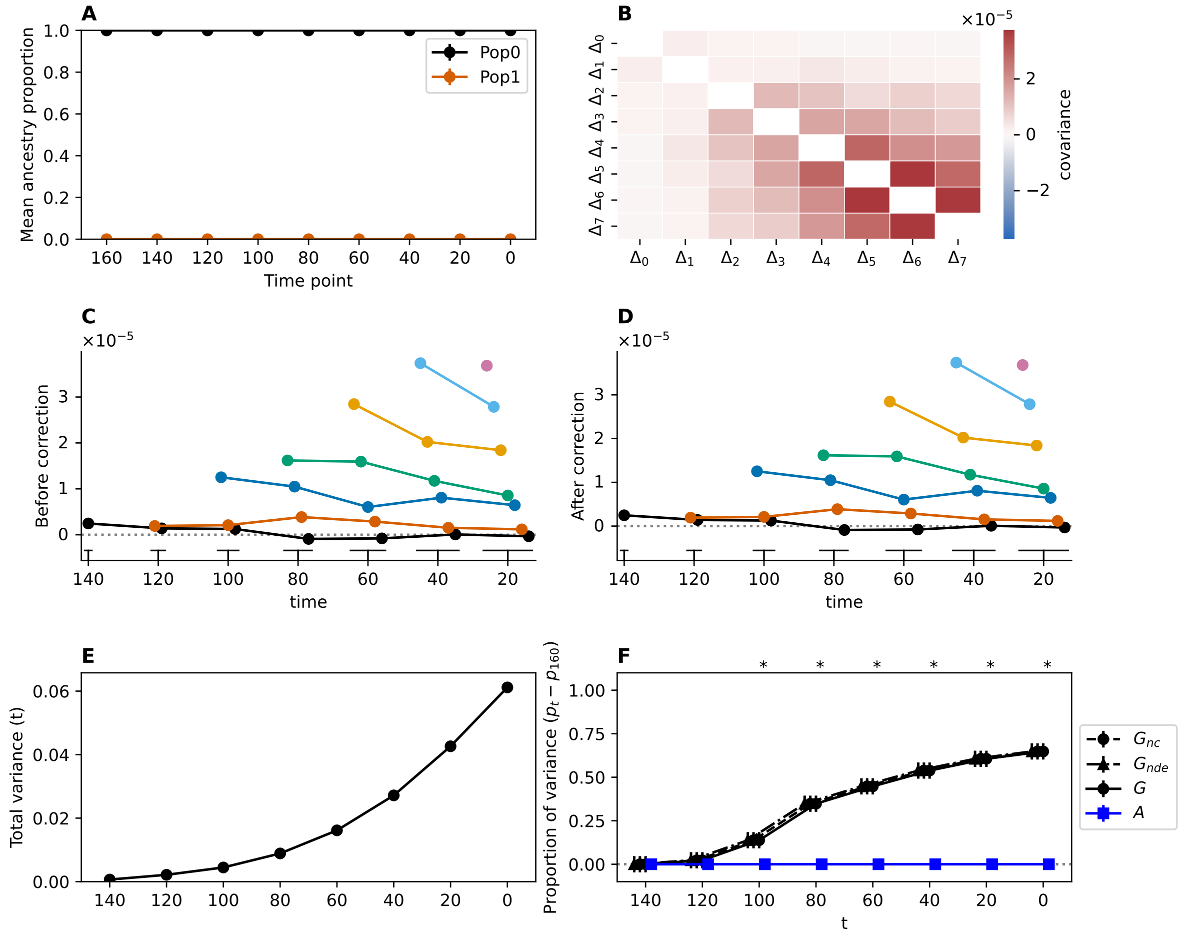


Figure S5: Demographic scenario with no gene flow (all fig. S1 pulses null), with selection as in fig. S3.

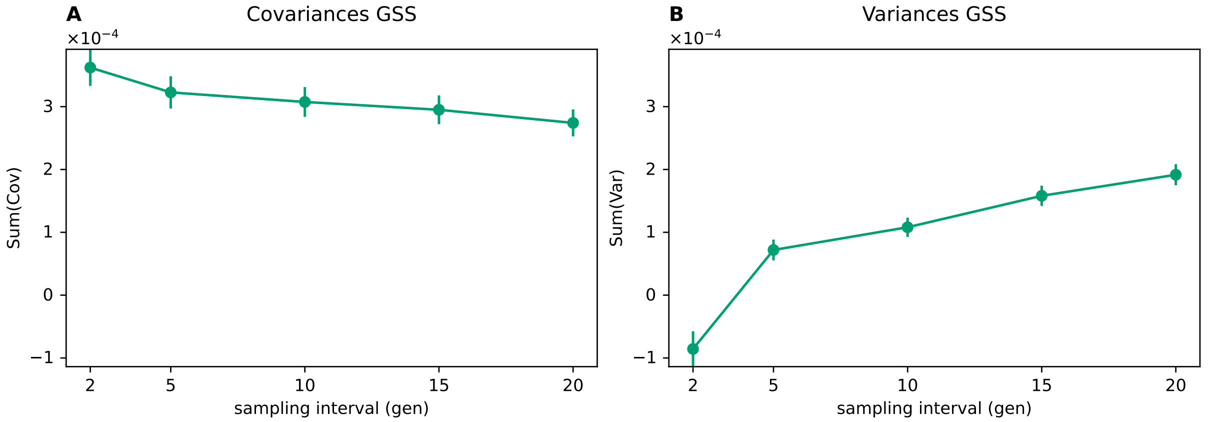


Figure S6: Total covariance (**A**) and variance (**B**) as a function of sampling time interval in the Gaussian stabilizing selection model (GSS) at a recombination rate of $r = 2 \times 10^{-8}$. This figure is a subset of fig. 4 at the same recombination rate as the main simulations to show the variation on a smaller y-axis range.

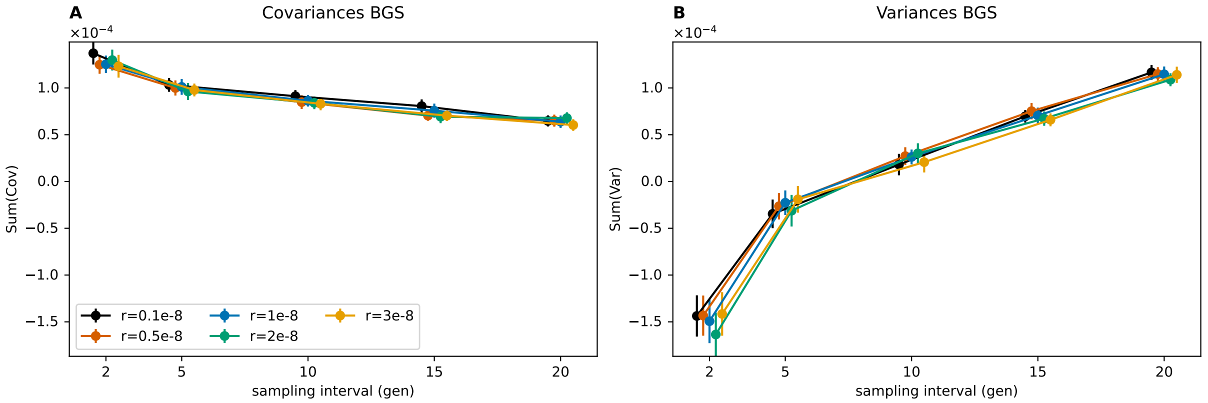


Figure S7: Total covariance (**A**) and variance (**B**) as a function of sampling time interval in the background selection model (BGS) with variable recombination rates (color coded). This figure is similar to fig. 4.

### Ancient DNA datasets

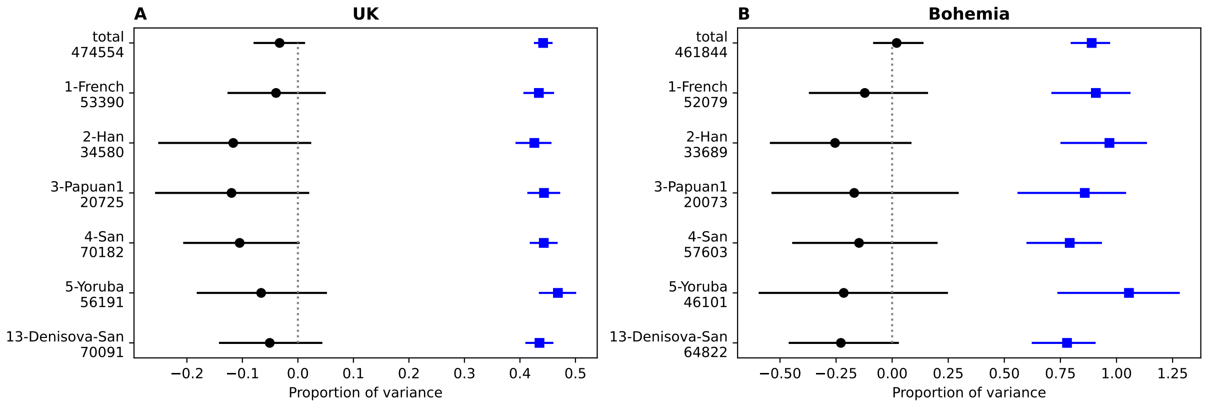


Figure S8: Investigation of the role of SNP ascertainment on the proportion of variance in the two analyzed datasets from UK (A) and Bohemia (B). Final time G and A proportions of variance, black circles and blue squares respectively, with 95 % block bootstrap CIs for the total set of SNPs (top row) and 6 subsets of SNPs that have been ascertained in different populations in the Human Origins SNP array design (Rohland et al., 2022, used column "24-ascertainment" from Supplement Data 7). The number of SNPs used in each case is indicated below the ascertained group label. Analysis was only performed for ascertained groups providing enough SNPs for the downstream computations to be relevant (on a total of 13 different groups).

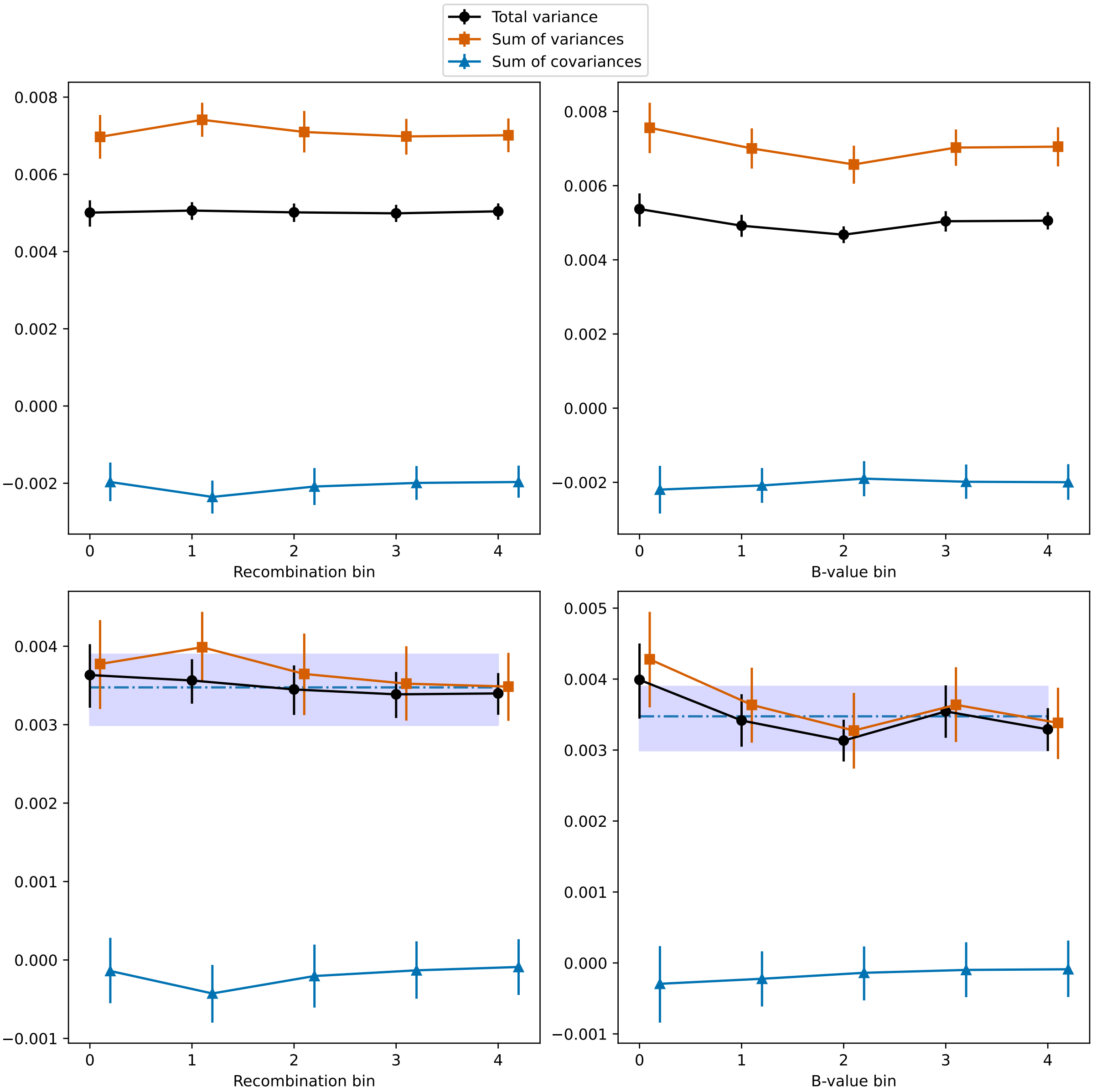


Figure S9: UK time transect total variance, sum of variances and sum of covariances binned for recombination (left panels) and B-values (right panels), either before (top panels) or after (bottom panels) admixture correction. The blue dash-dotted line and interval are the genome-wide mean and confidence interval for total variance computed with 1/5 of windows sampled on each bootstrap to be comparable to the binned values. Recombination bins: [0-0.079), [0.079-0.205), [0.205-0.513), [0.513-1.556), [1.556-658.284). B-value bins: [0.536-0.755), [0.755-0.849), [0.849-0.902), [0.902-0.944), [0.944-0.997).

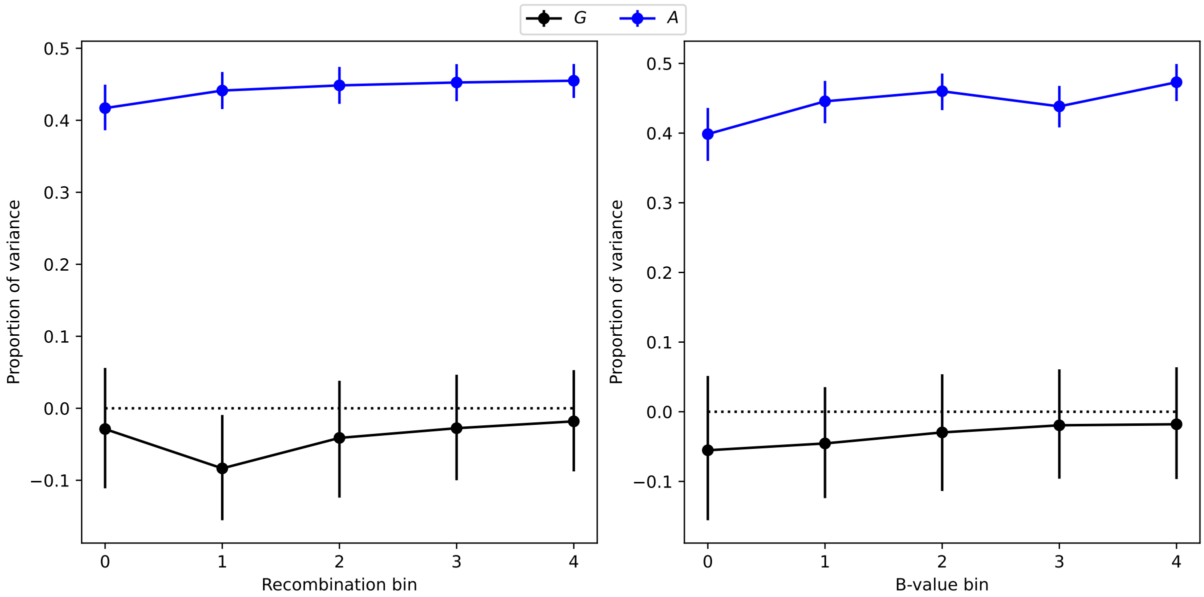


Figure S10: UK time transect $G$ and $A$ values. Recombination bins: [0-0.079), [0.079-0.205), [0.205-0.513), [0.513-1.556), [1.556-658.284). B-value bins: [0.536-0.755), [0.755-0.849), [0.849-0.902), [0.902-0.944), [0.944-0.997).

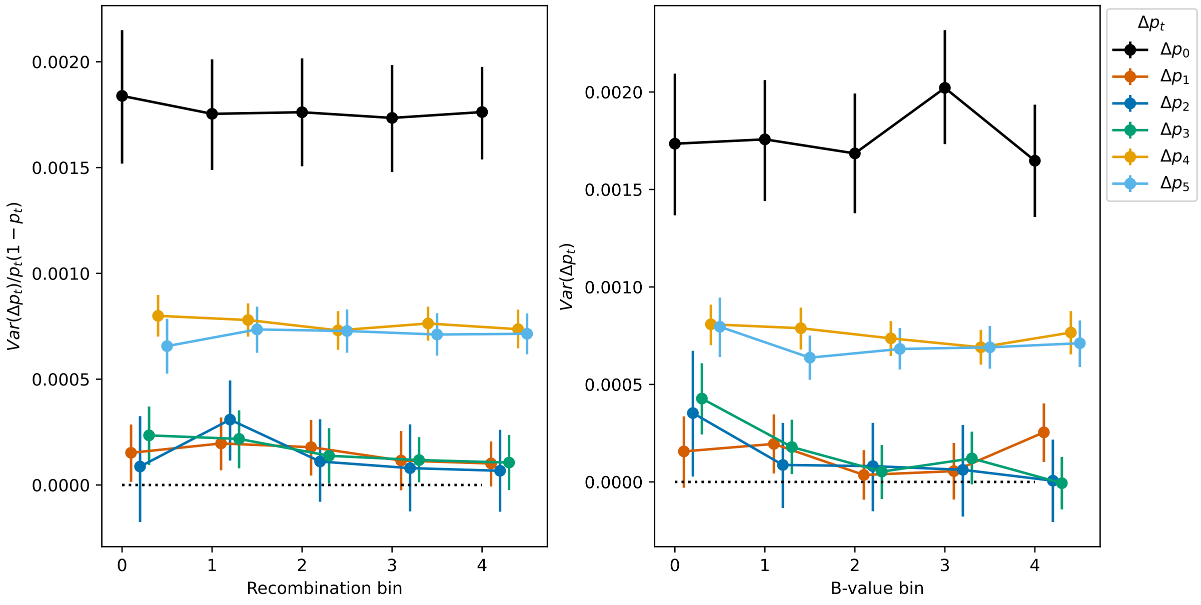


Figure S11: UK time transect variance of each time interval. Recombination bins: [0-0.079), [0.079-0.205), [0.205-0.513), [0.513-1.556), [1.556-658.284). B-value bins: [0.536-0.755), [0.755-0.849), [0.849-0.902), [0.902-0.944), [0.944-0.997).

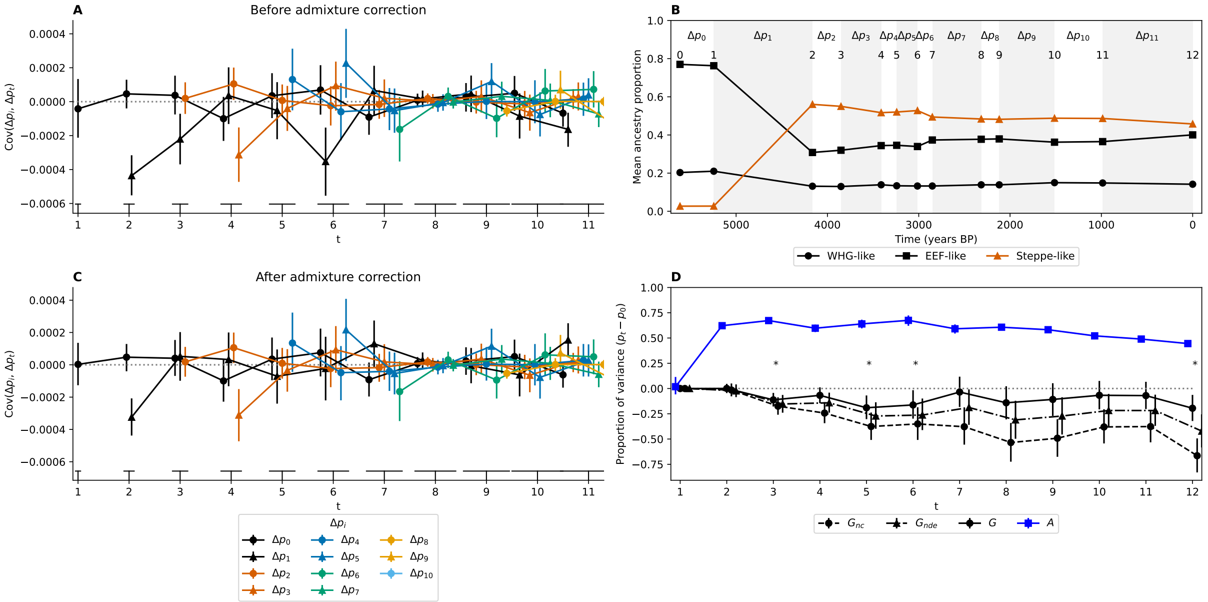


Figure S12: UK time series with time points split in two at the median sample age. All 95 % CIs are computed through a block bootstrap procedure and represented with vertical bars. **(A)** Covariance values before admixture correction. Each line corresponds to the covariance between a first time interval ($\Delta p_i$, color code) and a later time interval ($\Delta p_j$, x-axis). **(B)** Mean ancestry proportions from the three reference populations for each temporal sample. **(C)** Covariance values after admixture correction. **(D)** Proportion of the total variance, between the initial measured time (0) and time $t$, due to linked selection ($G_{nc}$ for non-corrected, $G_{nde}$ without drift error correction, and $G$) and to gene flow ($A$).

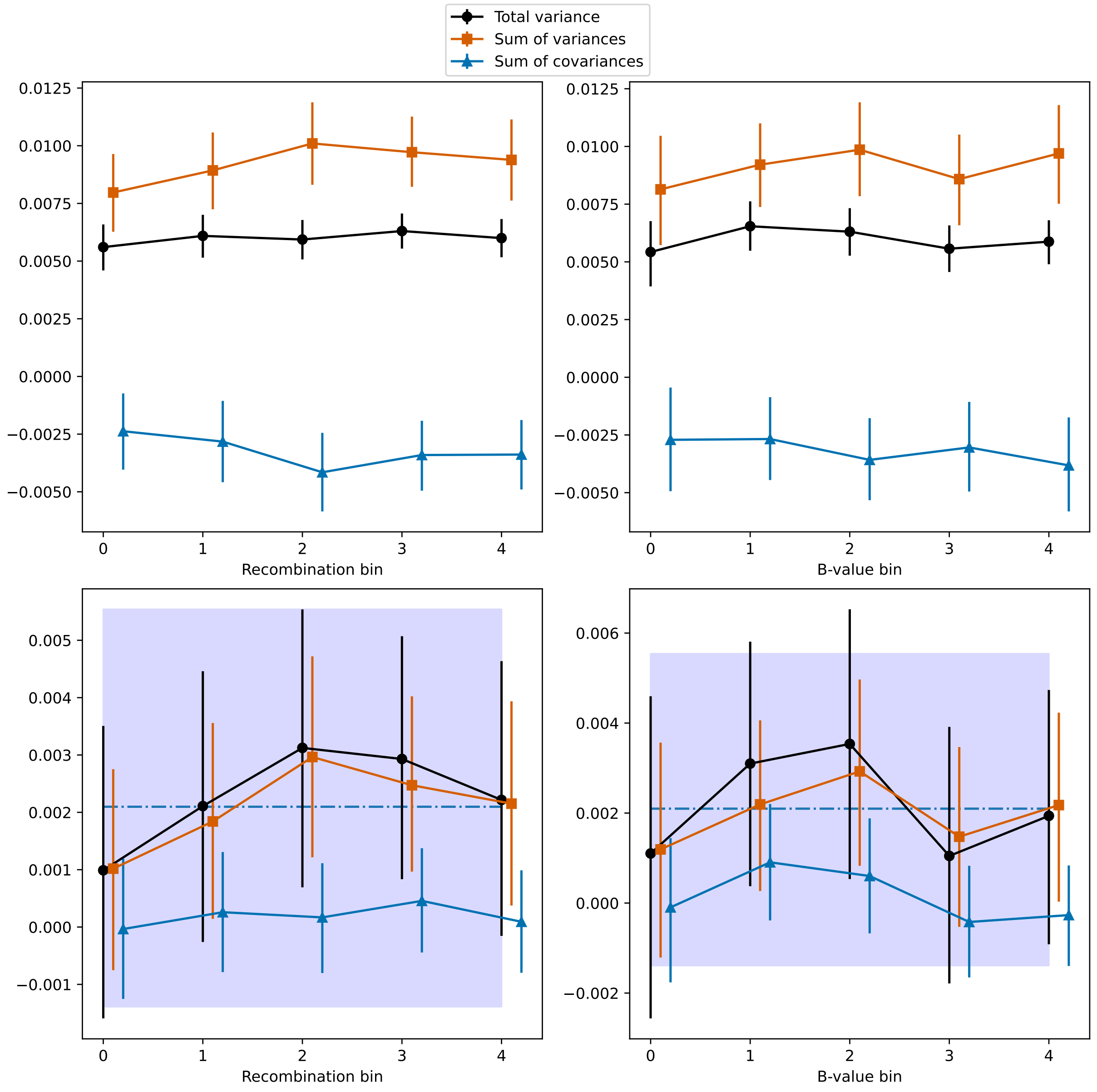


Figure S13: Bohemia time transect total variance, sum of variances and sum of covariances binned for recombination (left panels) and B-values (right panels), either before (top panels) or after (bottom panels) admixture correction. The blue dash-dotted line and interval are the genome-wide mean and confidence interval for total variance computed with 1/5 of windows sampled on each bootstrap to be comparable to the binned values. Recombination bins: [0-0.080), [0.080-0.208), [0.208-0.522), [0.522-1.573), [1.573-756.637). B-value bins: [0.536-0.757), [0.757-0.850), [0.850-0.902), [0.902-0.944), [0.944-0.997).

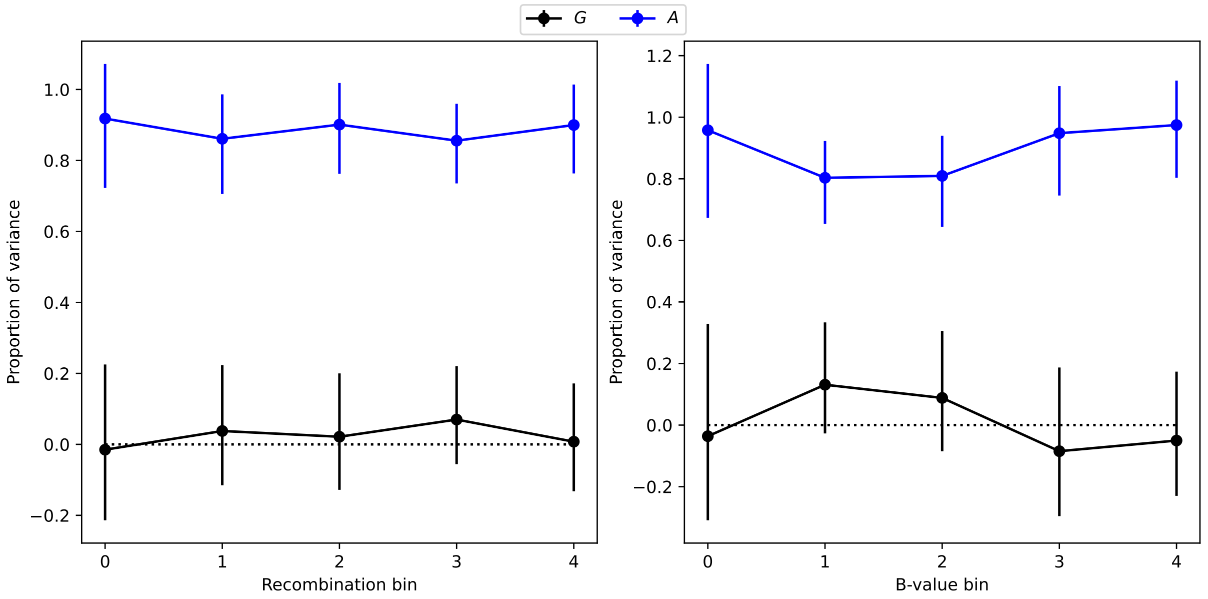


Figure S14: Bohemia time transect $G$ and $A$ values. Recombination bins: [0-0.080), [0.080-0.208), [0.208-0.522), [0.522-1.573), [1.573-756.637). B-value bins: [0.536-0.757), [0.757-0.850), [0.850-0.902), [0.902-0.944), [0.944-0.997).

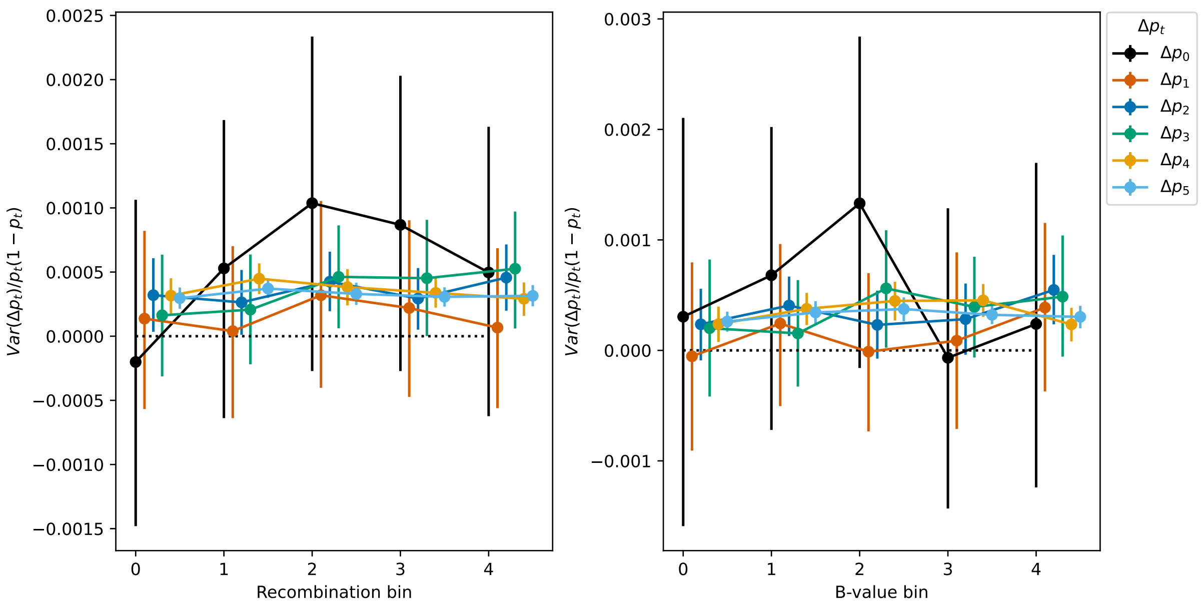


Figure S15: Bohemia time transect variance of each time interval. Recombination bins: [0-0.080), [0.080-0.208), [0.208-0.522), [0.522-1.573), [1.573-756.637). B-value bins: [0.536-0.757), [0.757-0.850), [0.850-0.902), [0.902-0.944), [0.944-0.997).

#### UK-like simulation

To give a sense of the expected neutral pattern observed in the UK time transect (Patterson et al., 2022), we simulated the admixture pulses inferred from the real data. We took as a base the realistic model developed by Allentoft et al., 2022 available in `stdpopsim`. The modified demography is presented in fig. S16 and the `demes` file is available in the github repository for simulations (https://github.com/alxsimon/admixcov_sims/blob/main/resources/AncientEurope_4A21_mod.yaml).

Similar to the main text simulations, we used `msprime` to create 100 replicates of neutral simulations using the Human chromosome 1 parameters of `stdpopsim`. Results are presented in fig. S17.

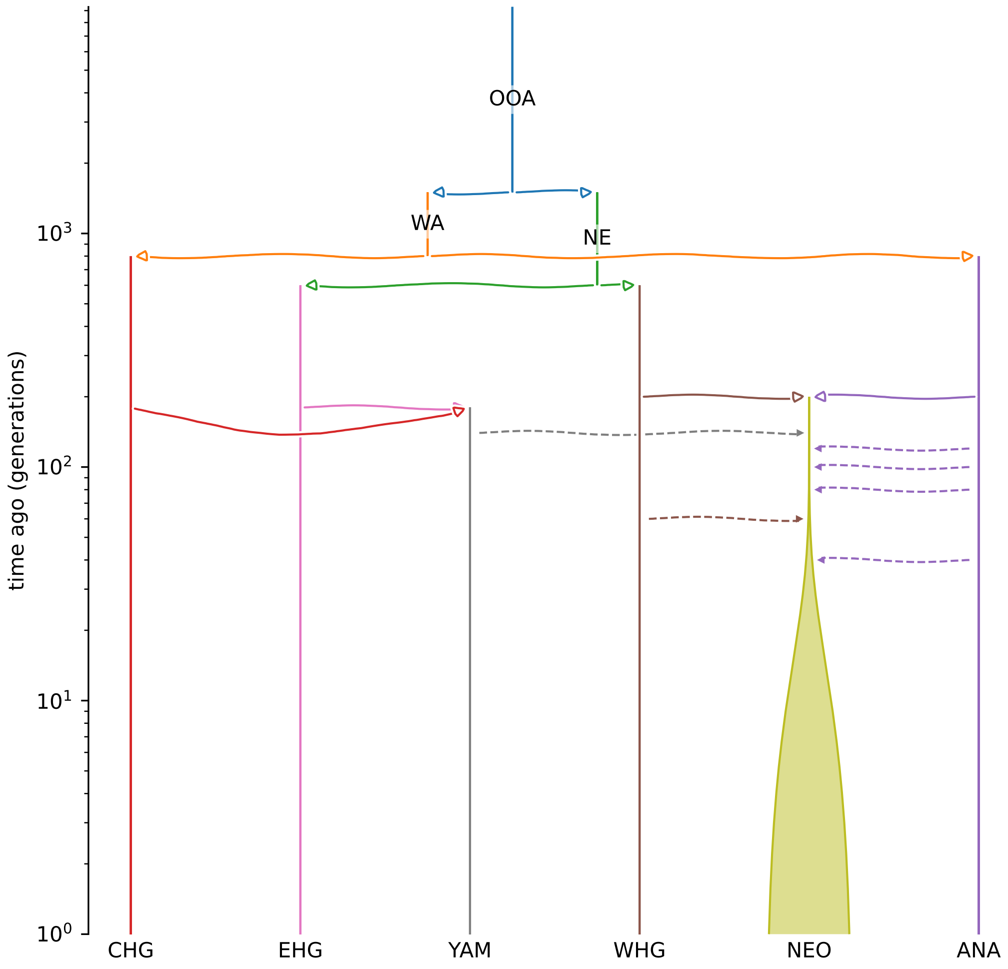


Figure S16: UK-like time transect simulation demography, modified from Allentoft et al., 2022's `stdpopsim` model. CHG: Caucasus Hunter Gatherers, EHG: Eastearn Hunter Gatherers, YAM: Yamnaya, WHG: Western Hunter Gatherers, NEO: Neolithic European population (and modern populations), ANA: Anatolia (Early farmers), WA: West Asia, NE: Northern Europe, OOA: Out of Africa population.

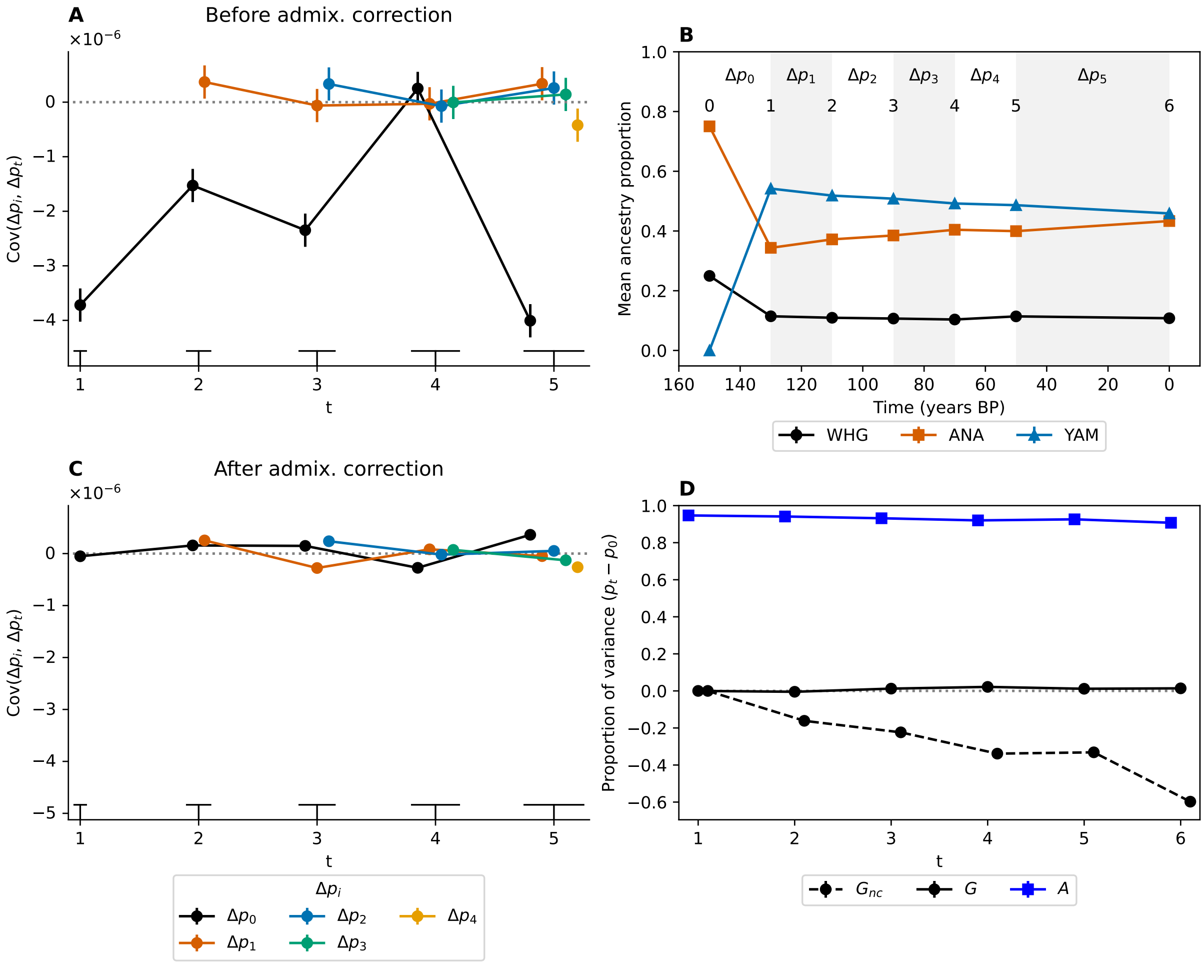


Figure S17: UK-like simulation results.
